# Cryo-EM Structures of the XPF-ERCC1 Endonuclease Reveal an Auto-Inhibited Conformation and the Basis for Activation

**DOI:** 10.1101/796524

**Authors:** Morgan Jones, Fabienne Beuron, Aaron Borg, Andrea Nans, Christopher Earl, David C. Briggs, Maureen Bowles, Edward P. Morris, Mark Linch, Neil Q. McDonald

**Affiliations:** Signalling and Structural Biology Laboratory, Francis Crick Institute, NW1 1AT, London, United Kingdom; Structural Electron Microscopy, The Institute of Cancer Research, SW7 3RP, London, United Kingdom; Mass Spectrometry Science Technology Platform, Francis Crick Institute, NW1 1AT, London, United Kingdom; Structural Biology of Cells and Viruses, Francis Crick Institute, NW1 1AT, London, United Kingdom; Department of Oncology, University College London Cancer Institute, WC1E 6AG, London, England, United Kingdom; Institute of Structural and Molecular Biology, Birkbeck College, Malet Street, London WC1E 7HX, United Kingdom

## Abstract

The structure-specific endonuclease XPF-ERCC1 participates in multiple DNA damage repair pathways including nucleotide excision repair (NER) and inter-strand crosslink repair (ICLR). How XPF-ERCC1 is catalytically activated by DNA junction substrates is not currently understood. We report cryo-electron microscopy structures of both DNA-free and DNA-bound human XPF-ERCC1. DNA-free XPF-ERCC1 adopts an auto-inhibited conformation in which the XPF helical domain masks ERCC1 DNA-binding elements and restricts access to the XPF catalytic site. Binding of a model DNA junction separates the XPF helical and ERCC1 (HhH)_2_ domains, promoting activation. Using these structural data, we propose a model for a 5’-NER incision complex involving XPF-ERCC1-XPA and a DNA junction substrate. Structure-function data suggest xeroderma pigmentosum patient mutations often compromise the structural integrity of XPF-ERCC1. Fanconi anaemia patient mutations often display substantial in-vitro activity but are resistant to activation by ICLR recruitment factor SLX4. Our data provide insights into XPF-ERCC1 architecture and catalytic activation.

## Introduction

Structure-specific endonucleases (SSEs) are found in all branches of life and play crucial roles in genome repair, replication and recombination^1^. These endonucleases act on similar DNA structures with defined polarity but use different catalytic mechanisms. The structurally-related XPF/MUS81 family are an important group of human 3’-nucleases that associate to form two active endonuclease heterodimers (XPF-ERCC1 and MUS81-EME1) and a pseudo-nuclease (FANCM-FAAP24)^2^. XPF-ERCC1 recognises double-stranded/ single-stranded (ds/ss) DNA junctions which have a 3’-ssDNA overhang, nicking the dsDNA backbone to produce a substrate for subsequent steps in repair path ways. XPF-ERCC1 activity is essential for removing helical DNA distortions arising from UV-induced damage and bulky adducts as part of the nucleotide excision repair (NER) pathway^3^. In this context XPF-ERCC1 nicks the damaged DNA strand 5’ of the lesion at the ds/ss junction of an NER repair bubble. It is also required for inter-strand crosslink repair (ICLR), some double-strand break repair processes, base excision repair, Holliday junction resolution, gene-conversion and telomere maintenance^4-10^. Mutations in XPF and ERCC1 genes are associated with genetic disorders exhibiting diverse phenotypes. These pathologies are caused by defects in the genome maintenance pathways that involve XPF-ERCC1 and include xeroderma pigmentosum (XP), Cockayne’s syndrome, Fanconi anaemia (FA), XPFE progeria and cerebro-oculo-facio-skeletal syndrome^11-15^. The genotype-phenotype correlations of XPF-ERCC1 driven diseases are still poorly understood.

XPF is the enzymatically active subunit of the heterodimeric endonuclease and is comprised of a helicase-like module (HLM) and a catalytic module (CM) (Fig. 1a). The XPF HLM is related to the superfamily 2 helicases, with two divergent RecA-like domains that flank an all α-helical domain^16^ (Fig. 1a). Both XPF RecA-like domains, termed RecA-like domain 1 (RecA1) and RecA-like domain 2 (RecA2) lack the residues necessary to bind and hydrolyse ATP^17,18^. Instead the HLM is required for full XPF activity and can bind to both the ICLR recruitment factor SLX4 and to ds/ssDNA structures^19,20^. The XPF CM consists of a nuclease domain containing a metal-dependent GDX_n_ERKX_3_D active site motif and a tandem helix-hairpin-helix termed an (HhH)_2_ domain^21^. The smaller ERCC1 subunit has no catalytic activity but is structurally related to the XPF CM, consisting of a nuclease-like domain (NLD) and its own dsDNA-binding (HhH)_2_ domain. Both ERCC1 domains heterodimerise with their equivalent domains in the XPF CM, forming discrete nuclease-NLD and 2x(HhH)_2_ functional units. As well as contributing to XPF stability, ERCC1 binds ds/ssDNA substrates and engages the XPA repair protein that is required for NER recruitment^22^. Currently there are no available structures of the XPF HLM or of any full-length XPF-Mus81 family members. By solving the structure of a near full-length human XPF-ERCC1 we have defined its overall architecture and uncovered a previously unreported auto-regulatory mechanism. We show XPF-ERCC1 adopts an auto-inhibited conformer in the absence of DNA in order to prevent promiscuous cleavage and provide structural evidence for initial steps in XPF-ERCC1 activation upon binding a DNA junction.

**Figure 1.**
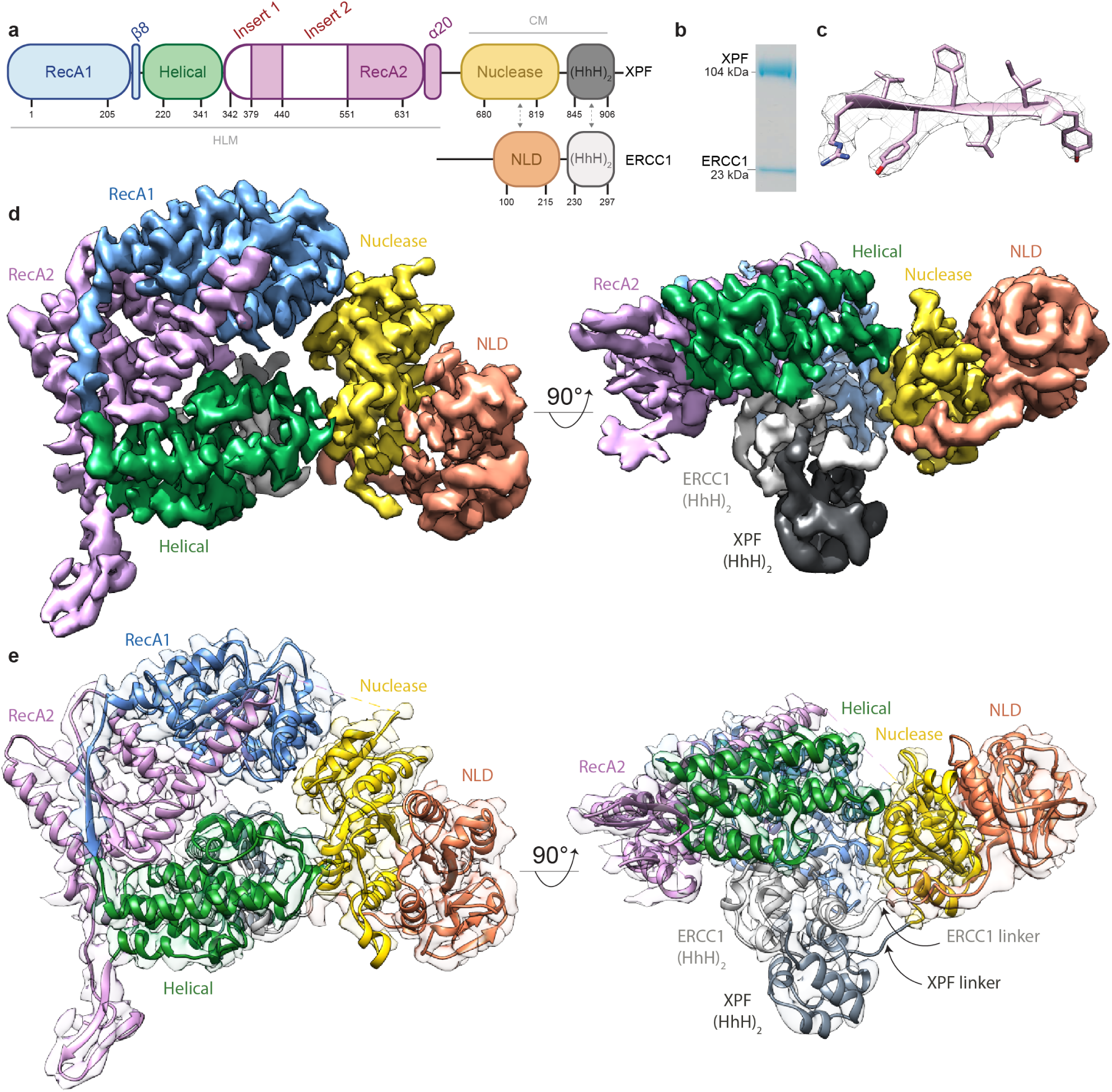
Structure of auto-inhibited human XPF-ERCC1 endonuclease. **a** Domain architecture of XPF-ERCC1 colour coded by domain. XPF: RecA1 (blue), Helical (green), RecA2 (purple), Nuclease (gold), (HhH)2 (dark grey). ERCC1: NLD (orange), (HhH)2 (light grey). Residue numbering indicates domain boundaries and dotted arrows indicate dimerisation interfaces. Two insert sequences within RecA2 are shown in white embellishing the RecA domain fold. Grey lines define the helicase-like module (HLM) and catalytic module (CM). **b** SDS-PAGE gel of purified recombinant XPF-ERCC1 used for cryo-EM studies. **c** Representative density of the cryo-EM map at 3.4 Å overlaid with a ß-strand and side-chains from the final model in pink. **d** Two orthogonal views of the composite XPF-ERCC1 cryo-EM map ranging from a global resolution of 3.6 – 4 Å, coloured by domain according to panel a. **e** Final XPF-ERCC1 atomic model coloured by domain according to panel a, displayed within the cryo-EM map density. The XPF nuclease-(HhH)2 and ERCC1 NLD-(HhH)2 domain linker density is apparent only at lower map thresholds.

## Results

### Structure determination of human XPF-ERCC1 endonuclease

A single particle cryo-electron microscopy (cryo-EM) density map of purified recombinant XPF-ERCC1 complex (128kDa) (Fig. 1b) was determined at a global resolution of 4.0 Å (Supplementary Fig. 4b) enabling the assignment of XPF-ERCC1 domain organisation. The map represents the single dominant conformer observed following 3D classification protocols (Supplementary Fig. 3) and exhibits clear secondary structure features throughout (Fig. 1c and 1d). Local resolution analysis indicated that the heterodimeric 2x(HhH)_2_ domain dimer exhibited some mobility, so signal subtraction of this domain was carried out followed by local refinement. This process improved the resolution of the resulting sub-volume to 3.6 Å (Supplementary Fig. 4d) which enabled de novo building, refinement and validation of an atomic model (Table 1). The locally-refined map shows clear side-chain density throughout with the local resolution ranging from 3.4 Å in the RecA1 and RecA2 domain cores (Fig. 1c) to 7 Å at the periphery of the ERCC1 NLD. Regions modelled as poly-alanine or omitted from the final structure are shown in Supplementary Table S1. There was no density recovered for the ERCC1 N-terminus, consistent with it being proteolytically cleaved (Supplementary Fig. 1b). Inspection of the angular distribution of assigned particle images during refinement, the 3DFSC curves and 3D flexibility analysis indicate that resolution differences were due to intrinsic flexibility rather than a lack of contributing particle images (Supplementary Fig. 4e-g).

**Table 1.** Cryo-EM statistics for XPF-ERCC1 structures and associated maps.

| Data Collection and Processing | XPF-ERCC1 | XPF-ERCC1<sub>DNA</sub> |
| --- | --- | --- |
| Voltage (kV) | 300 | 300 |
| Electron exposure (e-/Å<sup>2</sup>) | 63 | 63 |
| Defocus range (µm) | 1-4 | 1-3 |
| Pixel size (Å) | 1.38 | 1.38 |
| Symmetry imposed | C1 | C1 |
| Initial Particle Images | 3,989,887 | 3,432,565 |
| Final Particle Images | 405,339 | 199,022 |
| Map Resolution | 3.6 (local refinement)<br>4.0 (globally refined) | 7.9 (globally refined)<br>7.1 (local refinement) |
| Map Resolution Range | 3.4 - 8 | 5.5-14 |
| **Refinement** |  |  |
| Initial Model | Cryo-SPARC ab-initio | Cryo-SPARC ab-initio |
| Model Resolution (Å) | 3.6 - 4 | 5.9 - 7.7 |
| FSC threshold | 0.143 | 0.143 |
| Map sharpening B factor (Å<sup>2</sup>) | -168 | -560 |
| Model Composition |  |  |
| -Non-hydrogen atoms | 7014 | 7341 |
| -Protein residues | 943 | 892 |
| B factor (Å<sup>2</sup>) | 163 | 530 |
| RMSZ-score |  |  |
| - Bond lengths (Å) | 0.50 | 0.42 |
| - Bond angles (°) | 0.63 | 0.50 |
| Validation |  |  |
| -MolProbity score | 1.88 | 2.9 |
| -All-atom Clashscore | 15 | 14 |
| Ramachandran plot |  |  |
| -Favoured (%) | 89.0 | 93.0 |
| -Allowed (%) | 11.0 | 7.0 |
| -Disallowed (%) | 0 | 0 |

### Overall architecture of human XPF-ERCC1 endonuclease

The cryo-EM structure of near full-length XPF-ERCC1 reveals a compact conformation with extensive interactions between the XPF HLM and CM modules (Fig. 2a). Overall, the HLM adopts a “C”-shape that has dimensions of approximately 70Å x 40Å x 60Å. The two RecA-like domains form a rigid platform and lack a nucleotide cleft between them characteristic of many ATP-driven helicases. Instead the two XPF RecA-like domains are linked through the intimate intertwining of secondary structural elements that extend beyond their globular portion. While RecA1 caps one edge of the HLM and engages the XPF nuclease domain in the CM, the helical domain caps the other HLM extremity and engages the CM and the dsDNA-binding ERCC1 (HhH)_2_ domain (Fig. 2a). In this manner both ends of the HLM engage crucial elements within the XPF CM and ERCC1, indicating a key regulatory role for the HLM. Interfaces observed in the XPF-ERCC1 structure were largely validated using XL-MS data (Fig. 2b) (Supplementary Table S2). We note that this method requires accessible lysine sidechains therefore predominantly hydrophobic interfaces, such as the RecA1-nuclease, did not return any crosslinks^23^.

**Figure 2.**
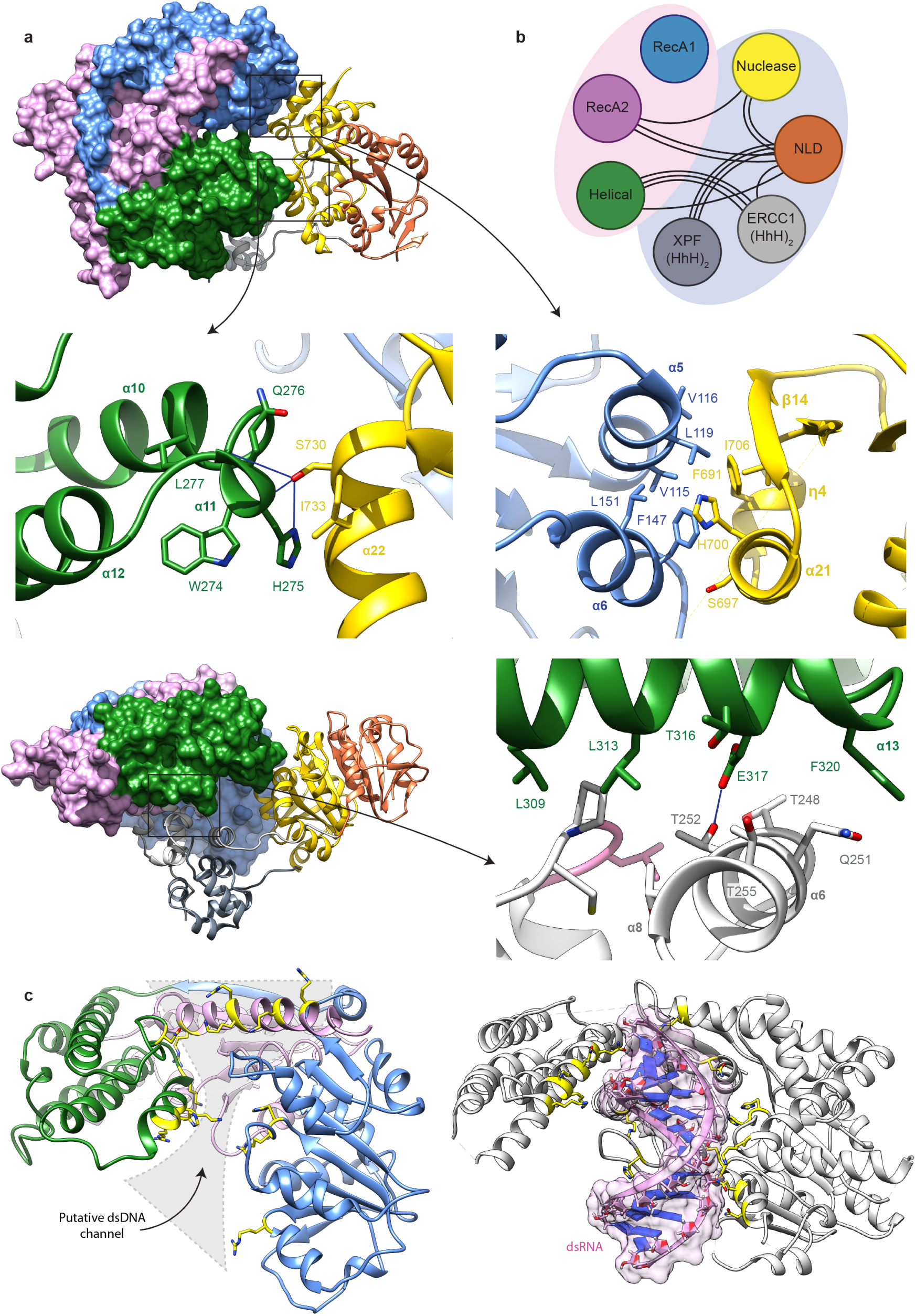
Architecture of the XPF helicase-like module and coupling with the catalytic module. **a** Upper panel, view of the XPF-ER-CC1 structure showing how the helicase-like module (HLM), displayed using surface rendering forms a “C”-shaped structure and contacts the XPF nuclease domain at two interfaces and the ERCC1 (HhH)2 domain, each shown as a cartoon. Domains are coloured according to the scheme used in Fig. 1a. Middle left panel, interaction between H275 in the XPF helical domain (green) and S730 in the XPF nuclease domain (gold). Blue lines indicate hydrogen bonds. Middle right panel, hydrophobic interaction interface between XPF RecA2 (blue) and nuclease (gold) domains. Lower right panel shows an orthogonal view and interaction between E317 in the XPF helical domain (green) and T252 in the ERCC1 (HhH)2 domain. **b** Cartoon schematic representing inter-domain crosslinks detected by mass spectrometry. Each black line indicates a single unique crosslink between residues in different domains. Domains within the pink ellipsoid form the HLM, whereas domains within the catalytic module and ERCC1 are within the pale blue ellipsoid. **c** left panel shows XPF HLM and basic residues lining a concave surface that could potentially bind to dsDNA. Right panel is an equivalent view of the helicase module from MDA5 (PDB code: 4GL2) bound to dsRNA through the same positively charged concave surface.

### Structure of the XPF helicase-like module (HLM)

The XPF HLM is typical of other helicase superfamily 2 (SF2) members with a RecA1-helical domain-RecA2 organisation, but with substantial inserts within RecA2. In the absence of ATP binding and hydrolysis motifs or a nucleotide binding cleft, RecA1-RecA2 are linked together through a predominantly polar interface (2007 Å^2^). Major interface contributions are made by intertwined secondary structural elements ß8 and α20 that form C-terminal extension to RecA1 and RecA2 respectively, as well as the XPF amino-terminus (Supplementary Fig. 5d). The main contacts centre on a π-ring stacking interaction between RecA1 domain Y71^XPF^ and RecA2 domain Y564^XPF^ at one interface edge (Supplementary Fig. 5c) and L39^XPF^ and I592^XPF^ on the other edge. Polar residues make up the remaining contacts with a small cavity. No protein expression was observed for a Y71A^XPF^ mutant (Table 2). The strand ß8 extends the smaller RecA2 four parallel ß-stranded sheet while α20 packs against the larger RecA1 seven-stranded parallel beta sheet (ß1–ß7). The observed structural rigidity of the RecA1-RecA2 unit is structurally homologous to equivalent domains in nucleosome-bound chromatin remodellers ISW1 and INO80^24,25^.

**Table 2.** Kinetic data for selected XPF-ERCC1 mutations.

| Protein | Mutant | Rationale | V<sub>max</sub><br>(fmol/min) | K<sub>m</sub><br>(nM) | k<sub>cat</sub><br>(s<sup>-1</sup>) | k<sub>cat</sub>/K<sub>m</sub><br>(nM<sup>-1</sup>min<sup>-1</sup>) |
| --- | --- | --- | --- | --- | --- | --- |
| XPF | WT |  | 104.7 ± 2.8 | 12.1 ± 1.4 | 20.9 ± 0.66 | 1.73 |
| XPF | L230R | FA | 65 ± 2.1 | 8.1 ± 1.1 | 13.0 ± 3.3 | 1.60 |
| XPF | L236R | FA/CS | 91.2 ± 2.5 | 10.8 ± 1.2 | 18.2 ± 0.5 | 1.69 |
| XPF | E239K | FA | 88.9 ± 4.7 | 12.4 ± 2.1 | 16.9 ± 0.7 | 1.36 |
| XPF | S786F | ICLR deficient | 80.9 ± 3.8 | 32.5 ± 1.5 | 7.9 ± 0.8 | 0.24 |
| XPF | 323-326 Δ | ICLR deficient | 120.2 ± 1.4 | 20.5 ± 0.8 | 24.2 ± 0.3 | 1.18 |
| XPF | E317A | Autoinhibition | 126.1 ± 28.2 | 5.1 | 13.5 ± 1.8 | 2.65 |
| XPF | W274A,<br>H275A | Autoinhibition | 193.4 ± 8.6 | 15.2 ± 2.4 | 38.7 ± 1.3 | 2.55 |
| XPF | R112A | DNA-binding | 24.5 ± 1.1 | 4.32 ± 0.63 | 5.6 ± 0.2 | 1.30 |
| XPF | 829-833 Δ | Autoinhibition linker disruption | 20.2 ± 0.5 | 1.9 ± 1.2 | 3.8 ± 0.2 | 2.00 |
| XPF | R589W | XP | 8.22 ± 2.6 | 25.9 ± 6.1 | 1.4 ± 0.9 | 0.05 |
| XPF | Q300A | Structural integrity | N/A | N/A | N/A | N/A |
| ERCC1 | L253A | Autoinhibition | 33.1 ± 2.7 | 28.7 ± 22.3 | 3.4 ± 0.3 | 0.12 |
| Protein | Mutant | Rationale | Effect of mutation on protein expression |  |  |  |
| XPF | L608P | XP | Soluble aggregates |  |  |  |
| XPF | T567A | XP | Soluble aggregates |  |  |  |
| XPF | Y71A | Structural integrity | No expression |  |  |  |
| XPF | R799W | XP | No expression |  |  |  |
| ERCC1 | T248A | Autoinhibition | No expression |  |  |  |
| ERCC1 | T252A | Autoinhibition | No expression |  |  |  |

XPF RecA2 has two large inserts with unknown functions (Fig. 1a). Insert one (residues 345-377) separates the helical and RecA2 domains and insert two (residues 441-550) interrupts the RecA2 fold. There is sufficient density in our map to trace the backbone of residues 345-362 and 366-377 from insert one projecting away from the body of the structure. However, no density was recovered for insert two, consistent with predictions that this region is likely intrinsically disordered in the absence of DNA. Cross-linking mass spectrometry (XL-MS) data identified a large number of intra-insert crosslinks within inserts one and two, consistent with these highly basic regions being flexible (Supplementary Table 1).

The XPF helical domain is an integral part of the HLM and folds as a five anti-parallel helical bundle. This domain packs tightly against RecA2 in a conformation stabilised by the interactions between D302^XPF^ and Q419^X-PF^, and Q300^XPF^ and S412^XPF^ (Supplementary Fig. 5b). The Q300A^XPF^ mutant significantly reduces XPF-ER-CC1 expression and increases aggregation (Table 2). Helix α17 (residues 426-440) also contributes to tethering the helical domain to RecA2. The observed position of the helical domain determines the orientation and angle of the extended RecA2 C-terminal α20 helix (Supplementary Fig. 5d), stabilising the HLM conformation through interaction between Q226^XPF^ and T614^XPF^.

The closest structural homologue identified by DALI protein structural comparison server^26^ is the helicase/ translocase MDA5 that binds dsRNA^26-28^ (rmsd of 4.1Å over 283 C-alpha atoms) (Fig. 2c and Supplementary Fig. 10b). Although comparisons with XPF HLM suggests that the closed, auto-inhibited conformation of the XPF HLM would not be blocked from binding dsDNA, we note that MDA5 binds to the major groove of a dsRNA using a concave surface lined with basic residues and sequences equivalent to the XPF RecA2 insert two spanning residues 441-550 (Fig. 2c). A similar positively charged concave surface is evident for XPF HLM and the insert two residues are disordered in the DNA-free structure but may become ordered in the presence of dsDNA.

### Auto-inhibitory elements within the helical domain regulate XPF-ERCC1 activity

The XPF HLM is coupled to the CM through contacts from RecA1 and the helical domain (Fig. 2a). RecA1 forms a substantial interface (1684 Å^2^) with the XPF nuclease domain involving aromatic and hydrophobic residues from RecA1 α5 and α6 helices and XPF nuclease domain η4 and α21 helices and ß14 strand (Fig. 2a). The hydrophobic nature of the contact suggests that anchoring of the HLM to the XPF nuclease domain through RecA1 forms a obligate part of the XPF-ERCC1 architecture.

The XPF helical domain forms a contact with the XPF nuclease domain that sterically prevents the ds/ss-DNA substrate from reaching the XPF active site (Fig. 2a). A key contact within this auto-inhibited conformation is between sidechains of H275^XPF^ and S730^XPF^ (Fig. 2a, middle panel), the latter sidechain also forms multiple hydrogen-bonds to the mainchain backbone of the helical domain. A H275A^XPF^, W274A^XPF^ double mutant, likely to disrupt this contact, displays a 1.5-fold increase in catalytic efficiency relative to the wild-type (Table 2). A second auto-inhibitory interface exists between the XPF helical domain and the ERCC1 (HhH)_2_ domain (Fig. 2a, lower panel). This interface is formed through predominantly polar contacts involving the highly conserved T248^ERCC1^ and T252^ERCC1^ residues, the latter of which forms a hydrogen bond with E317^X-PF^. Previous structural and biochemical data suggest that the ERCC1 (HhH)_2_ domain binds dsDNA through hairpin residues S244^ERCC1^-N246^ERCC1^ and G276^ER-CC1^-G278^ERCC1^ mainchain atoms^29,30^. These motifs are proximal to T248^ERCC1^ and T252^ERCC1^, and are not accessible in the DNA-free conformation of XPF^29^. The E317A^XPF^ mutant displays a 1.5-fold higher catalytic efficiency than the wild-type likely due to the disruption of this auto-inhibitory interaction (Table 2). Equally, shortening the connecting linker between the XPF nuclease-and (HhH)_2_ domain would be predicted to shift the 2x(HhH)_2_ unit towards the nuclease domain. Indeed, a similar effect is observed for a 829-833Δ^XPF^ mutant displaying a modest 1.2-fold increase in catalytic efficiency and a 7.5-fold tighter Km relative to wild-type.

### XPF catalytic module heterodimerises with ERCC1 through two distinct interfaces

ERCC1 is intimately coupled to the XPF CM through two obligate dimerisation surfaces at the equivalent domains of each molecule (Fig. 1e). The XPF nuclease domain uses a helix-strand-helix motif (α25-ß19-α26) to heterodimerise with the equivalent surface of the ERCC1 NLD (α3-ß8-α4) forming a kidney-shaped dimer with an extensive interaction interface (1684 Å2) (Supplementary Fig. 5a). The contact is predominantly hydrophobic and is flanked by three salt-bridges (Supplementary Fig. 5a). This interface uses equivalent elements to those mediating heterodimerisation of homologous domains from Mus81-Eme1 and FANCM-FAAP24 complexes^31,32^. The observed interaction contradicts a previous report arguing against heterodimerisation of the XPF nuclease domain-ERCC1 NLD unit^30^. We note that the XPF (HhH)_2_ domain hetero-dimerises with the ERCC1 (HhH)_2_ domain through predominantly hydrophobic contacts close to F851^XPF^ and F900^XPF^ as previously observed^30,33^. The (HhH)_2_ domain from XPF and ERCC1 are connected to their XPF nuclease domain/ERCC1 NLD domain through ordered linker sequences. There is sufficient density in our cryo-EM map to trace the backbone for both linkers (Fig. 1e). The ERCC1 linker makes unexpected interactions with the XPF nuclease domain via Y215^ER-CC1^ and D221^ERCC1^ (Fig. 3b). We note that Y215^ERCC1^ lies adjacent to S786^XPF^ suggesting the FA mutation S786F^XPF^ would disrupt this contact with ERCC1.

**Fig. 3.**
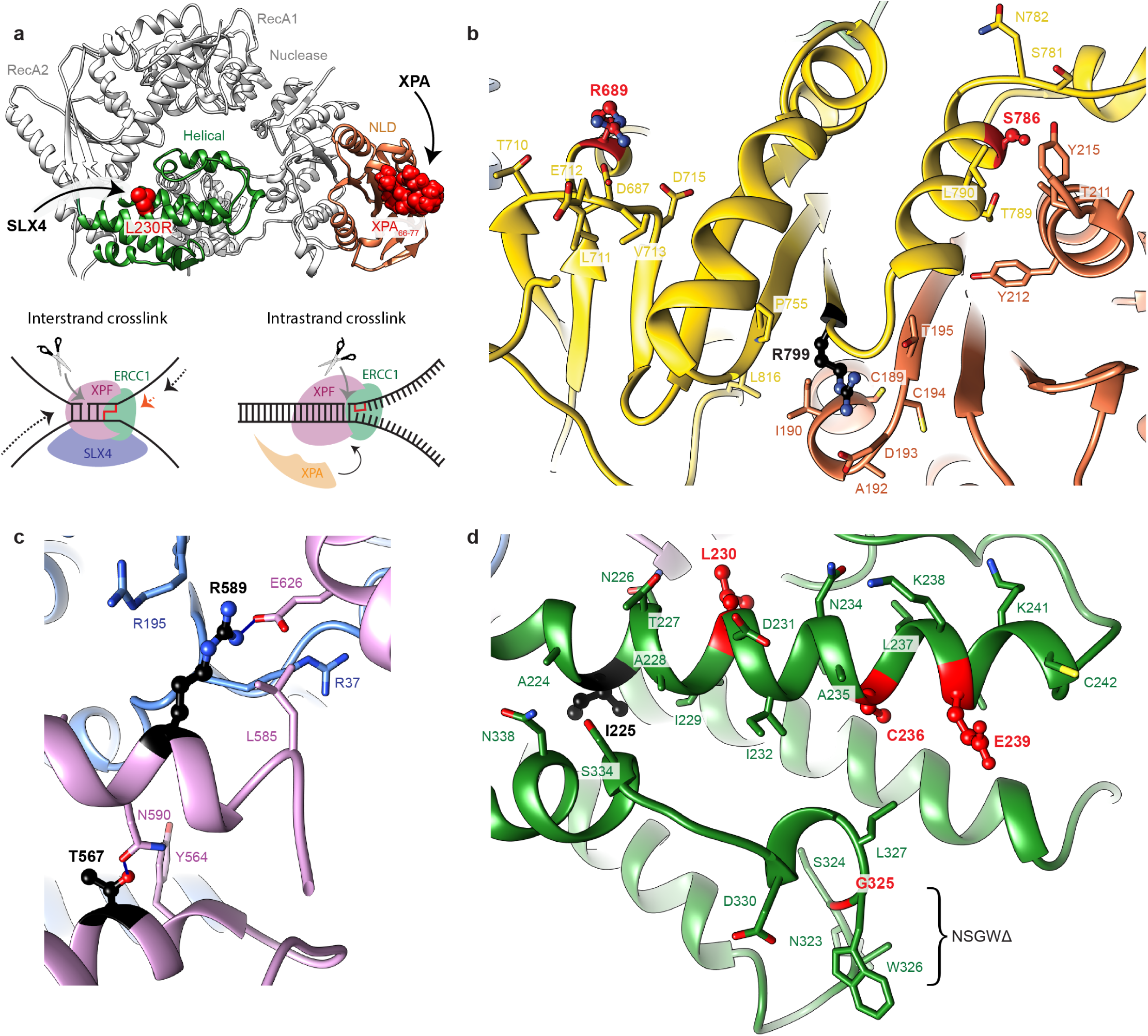
Mapping XPF-ERCC1 disease mutations and DNA repair pathway recruitment sites. **a** Ribbon model of XPF-ERCC1 highlighting the spatially distinct binding sites of XPA and SLX4. XPA binds to the ERCC1 NLD (coral) and SLX4 binds to the XPF helical domain (green). The XPA peptide (residues 66-77) atoms are displayed as red spheres (PDB: 2JNW). The key SLX4 binding residue L230 sidechain atoms are also displayed as red spheres. The DNA repair structures targeted by XPF-ERCC1 by recruitment through SLX4 (interstrand crosslink) or XPA (intrastrand crosslink) are shown. **b-d** The molecular environment of patient-derived disease mutations are indicated. Residues associated with patient mutations are displayed with ball and stick rendering and selected neighbouring residues are also shown as sticks. Residue colouring indicates the diseases associated with mutation of that residue: Red – Fanconi anaemia, Black – xeroderma pigmentosum. **b** Mutations in the XPF nuclease domain and ERCC1 NLD give rise to both Fanconi anaemia and XP. **c** XP associated mutations disrupt key structural contacts in the XPF RecA2 domain. **d** Fanconi anaemia associated mutations cluster in the XPF helical domain. The helical domain also contains the XP associated mutant, I225.

### Structural environment and differential impact of XP and FA patient mutations

Recruitment of XPF-ERCC1 into either NER or ICLR pathway complexes is dependent on interaction with partner proteins XPA or SLX4 at their respective damaged DNA structures (Fig. 3a). A previous study mapped the XPA-binding site to a cleft within the ERCC1 NLD^34^ (Fig. 3a). This interaction is spatially distinct from the proposed SLX4 site centred within the helical domain at L230^XPF 19^. Insights from disease-mutations have shown that repair pathway recruitment can be disrupted by separation-of-function (FA) or partial loss-of-function (XP) mutations, however the structural basis for this is unclear^35^.

With the availability of a three-dimensional XPF-ER-CC1 structure, it was possible to explore the location and structural environment of disease-causing mutations and correlate this with their impact on enzyme stability and catalytic activity. Patient-derived XP or FA associated mutations were characterised in-vitro using a previously reported fluorescence incision assay^20^. Mutations associated with XP mapped primarily to the XPF RecA2 domain and its inserts^15,36,37^. L608^X-PF^, R589^XPF^ and T567^XPF^ are located in the folded region of the RecA2 domain, with the latter two forming structurally important intra-domain contacts^36^ (Fig. 3c). Indeed, L608P^XPF^ and T567A^XPF^ mutant proteins formed soluble aggregates when expressed recombinantly, as measured by analytical size exclusion chromatography (data not shown), and an R589W^XPF^ mutant exhibited 35-fold reduction in catalytic efficiency (Table 2). The R799W^XPF^ XP mutation failed to express recombinantly and lies on the periphery of the heterodimeric nuclease-NLD interface with ERCC1 (Fig. 3b). These data, taken in the context of our structure, suggest the L608P^XPF^, T567A^XPF^, R589W^XPF^ and R799W^XPF^ XP disease mutants compromise XPF-ER-CC1 structural stability. I225^XPF^ in the helical domain (Fig. 3d) is also associated with XP but does not appear to contribute to XPF-ERCC1 structural stability^15^.

FA patients are proficient in NER but deficient in ICLR, indicating a separation of function^19,38^. FA point mutations within XPF including L230R^XPF^, C236R^XPF^ and G325E^XPF^ cluster in the XPF helical domain^11^ (Fig. 3d). These mutants, when expressed recombinantly, were found to have a similar level of endonuclease activity against a stem-loop substrate to wild-type XPF-ERCC1 (Table 2). SLX4 has been shown to stimulate XPF-ER-CC1 activity and recruit the endonuclease to ICLR sites ex-vivo^19^. In order to determine whether FA mutants display differences in activity in the presence of SLX4, an XPF-ERCC1-SLX4^NTD^ complex was assembled and its activity determined^39,40^ (Supplementary Fig. 11d). We found that the binding of SLX4^NTD^ to wild-type XPF-ER-CC1 stimulated its endonuclease activity, increasing the catalytic efficiency approximately 6-fold (Table 3). However, binding of SLX4^NTD^ to FA mutant 323-326Δ^XPF^ decreased the catalytic efficiency approximately 5.5-fold relative to wild-type XPF-ERCC1 in the absence of SLX4 (Table 3), suggesting that a subset of FA mutants are insensitive to SLX4 activation. FA mutant L230R^XPF^ was unable to bind SLX4^NTD^ (data not shown). This is consistent with a previous study that used full-length SLX4, indicating that it forms a key determinant of the SLX4 binding site^19^. Mutations that disrupt the SLX4 binding site of XPF-ERCC1 are likely to underpin FA disease by preventing recruitment to ICLR^41^.

**Table 3.**
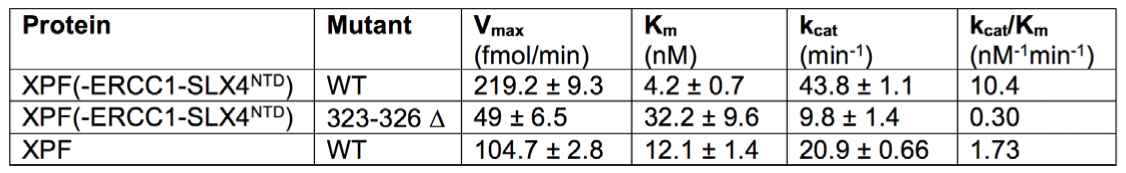
Activation of XPF-ERCC1 by SLX4^NTD^ and disruption by mutation.

### Evidence for XPF-ERCC1 conformational activation driven by DNA junction binding

The auto-inhibitory interactions formed by the XPF helical domain were hypothesised to be released following XPF-ERCC1 DNA junction engagement and prior to the incision reaction. To probe the nature of such potential conformational changes, we assembled a complex of XPF-ERCC1 bound to a DNA stem-loop substrate (10-duplex 20-T single-strand stem–loop) that we previously showed presents a single incision site to XPF-ER-CC^120^. Using an electrophoretic mobility shift assay (EMSA) we observed 1:1:1 stoichiometric binding of the stem-loop DNA to XPF-ERCC1 (Supplementary Fig. 6).

This sample was used for cryo-EM data collection leading to a single particle cryo-EM density map at a global resolution of 7.7 Å. This reconstruction enabled the placement of all XPF-ERCC1 domains using the DNA-free structure as an initial template (Fig. 4a). Signal subtraction of the dimeric 2x(HhH)_2_ domain and DNA density, followed by local refinement, improved the resolution of the resulting sub-volume to 5.9 Å (Supplementary Fig. 9c; Table 1). Comparison of both maps enabled the identification of the duplex portion of the stem loop based on the density for two strands and a concave surface of a dsDNA major groove (Fig. 4b). This enabled a DNA-bound XPF-ERCC1 structure to be built that revealed substantial conformational rearrangements uncoupling the XPF HLM from the CM. The greater conformational variability observed in the DNA-bound structure, compared to the DNA-free complex, is likely to have limited the resolution and may reflect the absence of stabilising recruitment factors or the need for longer DNA junction substrates.

The most striking changes to the XPF-ERCC1 structure following DNA binding involve the XPF helical and 2x(HhH)_2_ domains, consistent with an opening of the HLM and stem-loop engagement by the CM (Fig. 4c). The RecA1-RecA2 unit remains structurally rigid, with high-resolution features present in 2D class averages, whereas the rest of the complex increased in flexibility (Supplementary Fig. 7b). The interface between the XPF RecA1 and nuclease domains also remains intact. The resolution ranges from 7 Å in the core of the RecA1-RecA2 unit to 12 Å in the ERCC1 NLD (Supplementary Fig. 9a). Additional density is apparent adjacent to the RecA2 ß-sheet and could represent part of the missing insert two, analogous to a dsRNA-binding region of MDA5.

**Fig. 4.**
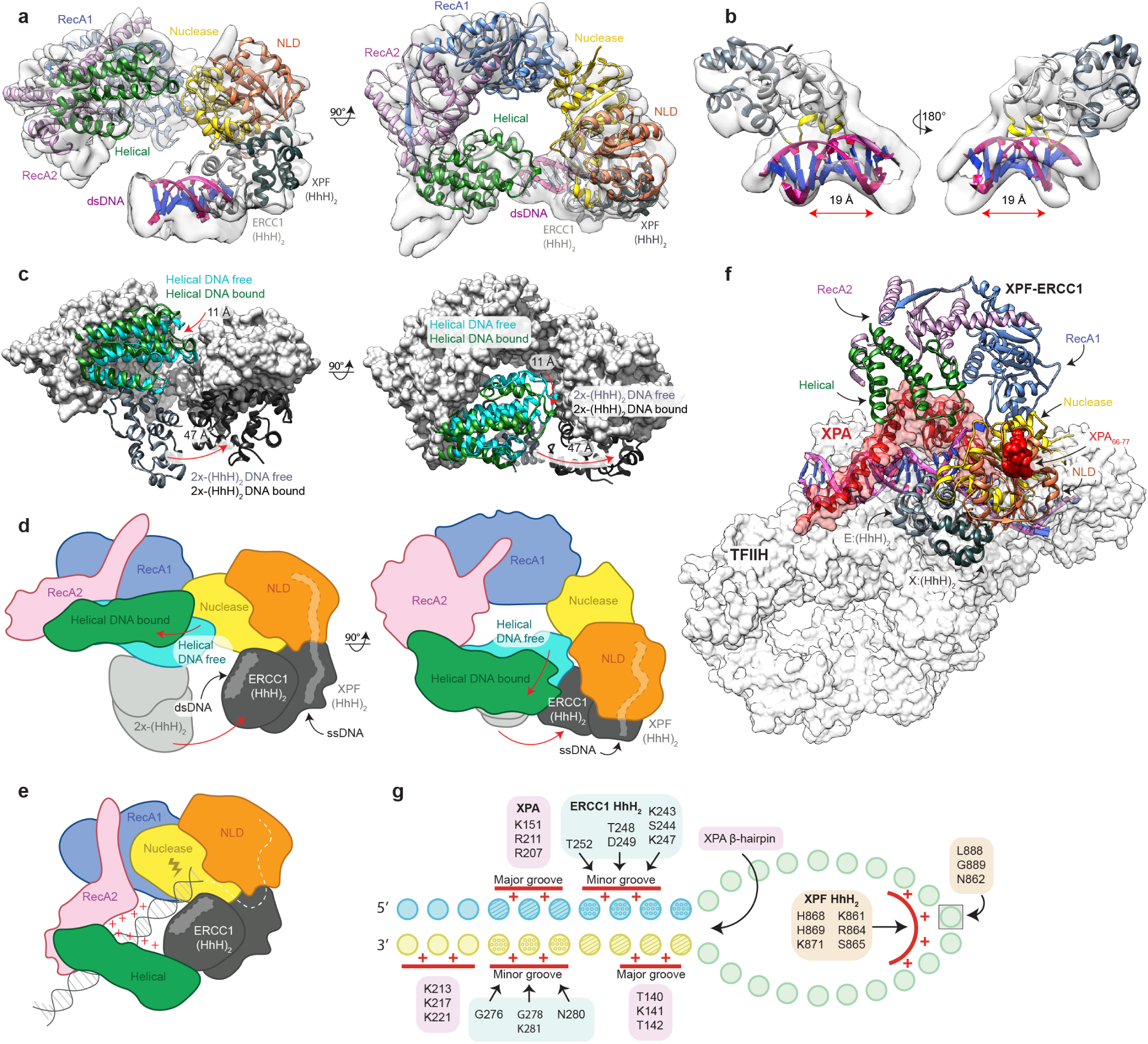
Evidence for conformational activation of XPF-ERCC1 by a DNA junction substrate. **a** Two orthogonal views of stem-loop DNA-bound XPF-ERCC1 coloured by domain according to Fig. 1a. Ribbon model placed within cryo-EM map density of composite map ranging from 5.9 – 7.7 Å global resolution. **b** Two views of segmented map density for the XPF-ERCC1 2x(HhH)2-dsDNA unit. XPF (HhH)2 domain grey, ERCC1 (HhH)2 domain white, dsDNA-binding hairpin residues of ERCC1 are highlighted in yellow, dsDNA purple and blue. Map density for the dsDNA shows an exposed major groove, measuring ∼19 Å across, and a buried minor groove engaged by the ERCC1 (HhH)2. **c** Conformational rearrangements required for DNA binding to XPF-ERCC1. Stem-loop DNA-bound XPF-ERCC1 model displayed as grey surface rendering. XPF in the absence of DNA (cyan), XPF helical domain in the presence of DNA (green), ERCC1 (HhH)2 domain in the presence of DNA (white), XPF (HhH)2 domain in the presence of DNA (black). Distances of domain rearrangements displayed in Å. **d** Cartoon of XPF-ERCC1 domain rearrangements upon binding stem-loop DNA. Red arrows indicate direction of domain rearrangements from DNA-free to DNA-bound. **e** Cartoon schematic of hypothetical, fully activated XPF-ERCC1 in which the major groove upstream of the ERCC1 HhH2 domain is engaged by the concave surface of the XPF HLM which feeds the DNA substrate into the active site. **f** Model for XPF-ERCC1 binding to ds/ssDNA-bound TFIIH-XPA. Stem-loop DNA-bound XPF-ERCC1 model docked onto the TFIIH-XPA-splayed arm DNA structure (PDB code 6RO4) through the exposed dsDNA minor groove at the ss/dsDNA boundary. XPF-ERCC1 model coloured according to domain as in Fig. 1a, DNA (purple), XPA (red) displayed with transparent surface rendering. TFIIH displayed in grey with opaque surface rendering. **g** Schematic of XPF-ERCC1 and XPA interactions with a stem-loop DNA substrate. XPA interaction elements displayed in pink, ERCC1 interactions displayed in blue, XPF interactions displayed in orange. There is a notable complementarity of XPA and XPF-ERCC1 binding to the splayed arm DNA and no mutually overlapping contacts consistent with a synergistic interaction.

Comparison with the DNA-free structure reveals that the XPF helical domain pivots by approximately 15°, rotating ∼11 Å away from the nuclease domain (Fig. 4c and Supplementary Fig. 10a). This conformational change breaks the autoinhibitory contact formed between H275^XPF^ and S730^XPF^. The positioning of the XPF α17 helix acts as a steric brake, preventing the HLM from adopting an more open conformation capable of binding dsDNA^27^ (Supplementary Fig. 10b). The most dramatic difference in the DNA-bound conformer is the displacement of the dimeric 2x(HhH)_2_ domain by 47 Å, detaching from the XPF helical domain to contact the XPF nuclease - ERCC1 NLD unit (Fig. 4c) similar to DNA-bound structures of XPF/Mus81 family members^29,31,32^ (Supplementary Fig. 10c).

The ERCC1 (HhH)_2_ domain engages the dsDNA through the minor groove through hairpin residues S244^ERCC1^-N246^ERCC1^ and G276^ERCC1^-G278^ERCC1^ main-chain atoms, similar to a previous structure of A. pernix XPF bound to dsDNA^29^. In this configuration the dimeric 2x(HhH)_2_ domain lies proximal to the ERCC1 NLD domain, pairing up both known ssDNA-binding elements of the endonuclease^29,31,32,42^ (ERCC1 NLD and XPF (HhH)_2_ domain) (Fig. 4d). Others have proposed that XPF-ERCC1 2x(HhH)_2_ domain is sufficient to recognise ds/ssDNA junctions^42^, however the precise arrangement of multiple ssDNA and dsDNA domains required for DNA junction recognition remains to be determined. The final DNA-bound model lacks the single-stranded portion of the stem loop and places the scissile phosphodiester bond approximately 15 Å from the XPF active site motif (residues 725-727). We interpret the DNA-bound structure as showing important features of an initial step towards full DNA junction recognition prior to the incision reaction. The low resolution of the DNA component within the cryo-EM map (approximately 9 Å) suggests that the dimeric 2x(HhH)_2_-DNA complex can adopt multiple conformers. Equally, the accessibility of the dsDNA major groove opposite to the 2x(HhH)_2_ minor groove interaction could be re-oriented towards the positively charged concave surface within the XPF HLM, analogous to how MDA5 recognises dsRNA (Fig. 4e).

### Towards a complete model for nucleotide excision repair 5’-incision complex

While this manuscript was in preparation, the structure of a ds/ssDNA-bound TFIIH-XPA (PDB code: 6RO4) was published representing a 5’-NER pre-incision complex that XPF-ERCC1 is known to engage^43^. We therefore sought to build a full model for an NER 5’-incision complex, by docking our DNA-bound XPF-ER-CC1 conformer onto the structure of ds/ssDNA-bound TFIIH-XPA. To do this we superimposed the ERCC1 (HhH)_2_ domain-dsDNA complex onto the exposed DNA minor groove at the ds-ssDNA junction (Fig. 4f) noting a remarkable non-overlapping complementarity in DNA binding with XPA. ERCC1 engaged precisely the available DNA elements that were not engaged by XPA (Fig. 4g). The endonuclease activity of XPF-ERCC1 nicks two nucleotides into the duplex portion of a model stem-loop^20^. As XPF-ERCC1 lacks an obvious “wedge” to separate dsDNA near discontinuities, but has both dsDNA and ssDNA binding functions, it is possible that XPA and SLX4 serve in this capacity in NER and ICLR respectively. Consistent with this XPA utilises a ß-hairpin element to separate dsDNA individual strands and demarcates the 5′-edge of the repair bubble^43^.

The 5’-incision complex model also anticipates several extensive interfaces between the XPF-ERCC1 and TFIIH-XPA-DNA structures and contained surprisingly few steric clashes, all of which were within the flexible XPA loop region (residues 104-131) (Fig. 4f). In this model, the dimeric 2x(HhH)_2_ domain lies adjacent to the TFIIH subunit XPB and DNA whilst the XPF nuclease-ERCC1 NLD dimer is positioned close to XPD, XPA and DNA. The highly basic and flexible RecA2 insert one (residues 345-377) is oriented to interact with either the extended XPA helix or dsDNA. The flexible regions of the XPF-ERCC1-DNA map correspond to those domains proposed to make extensive TFIIH-XPA-DNA interactions. It is therefore likely that these interfaces become stabilised upon TFIIH-XPA-DNA binding. Further experiments to validate these contacts by XL-MS are required.

## Discussion

The structural and functional studies described in this report provide new insights into XPF-ERCC1 architecture, regulation and activation. This XPF-ERCC1 endonuclease catalyses the first irreversible step in NER repair by nicking the 5’-edge of the repair bubble structure on the damaged strand. The structure of DNA-free XPF-ERCC1 reveals how the heterodimer is auto-inhibited by blocking both DNA binding and active site access through contacts with the XPF helical domain. This structure reveals inter-domain interfaces not previously described and rationalises our previous report that the HLM impacts on endonuclease activity and substrate interaction^20^. It also confirms the presence of a heterodimeric interface between the XPF nuclease and ERCC1 NLD as described for other family members^29,31,32^.

This study provides evidence linking conformational activation of XPF-ERCC1 through its recruitment by partner proteins to DNA junction sites prepared for either NER or ICLR pathways (Fig. 3a). Mapping the XPA interaction site within ERCC1^34^ and the SLX4 site within XPF helical domain reveals spatial separation of each recruitment partner site in the auto-inhibited state. It suggests the critical binding determinants are non-overlapping, but full structures of XPF-ERCC1 with SLX4 or XPA combined with competition binding studies are required to prove this. XPF-ERCC1 activation by SLX4 is disrupted by some FA mutations that map to the helical domain, in agreement with previous in-vivo work^19,38^. Given its proposed regulatory role, the helical domain may be repositioned on binding SLX4 to stimulate activity^39,40^. In contrast, XP associated mutations were found to generally reduce endonuclease activity in-vitro towards an NER substrate by destabilising the complex whereas FA mutants exhibited activity similar to wild-type.

XPF-ERCC1 cryo-EM structures described here reveal how engagement with a DNA junction substrate is able to disengage the XPF helical domain from the XPF CM and release the heterodimeric 2x(HhH)_2_ domain. The released 2x(HhH)_2_ domain is then able to engage a minor groove in the DNA duplex adjacent to the DNA ds/ss junction and packs against the XPF nuclease - ERCC1 NLD dimer, as observed for structures of Mus81-Eme1 and A. pernix XPF. In-silico docking of the DNA-bound conformation of XPF-ER-CC1 onto the recent TFIIH-XPA-DNA structure permits the assembly of a model for a NER 5’-incision complex. The model is consistent with the ERCC1 (HhH)_2_ hairpins engaging the minor groove and XPA the major groove of a DNA substrate. This positions the XPF hairpins to bind ssDNA and the nuclease domain for cleavage of substrates with the correct polarity.

The structures described here do not reveal the full basis for DNA junction recognition or the extent of conformational flexing required to place the scissile bond proximal to the XPF catalytic centre. We speculate that the similarities between XPF HLM and the MDA5 helicase point to a concave surface that could engage the major groove of the dsDNA element of a DNA junction to promote movement of the ds-ssDNA discontinuity into the XPF catalytic site. Evidently further high-resolution structures are required with longer DNA substrates and recruitment partner complexes in order to fully understand how the scissile phosphodiester bond is presented to the XPF catalytic site and the extent of the conformational alterations required.

There is a pressing need to explore opportunities for chemical inhibition of XPF-ERCC1 in view of the XPF synthetic lethality with the DNA translocase FANCM, particularly relevant to FANCM-deficient tumours^44^ and also with PARP inhibition^45^. The availability of an atomic structure for human XPF-ERCC1 described here will encourage such efforts to develop new precision medicines as well as to overcome cancer chemoresistance ^46^.

## Methods

### XPF-ERCC1 expression, purification and complex assembly

All reagents purchased from Sigma-Aldrich unless otherwise stated. A pFastBac Dual vector containing full length, wild type, human XPF (NCBI reference sequence: NM_005236.2) and ERCC1 (NCBI reference sequence: NM_001166049.2) cDNA was modified to include a C-terminal ERCC1 dual strep-tag using restriction enzyme cloning. This plasmid was transformed into competent DH10BAC E. coli cells (Thermo-Fisher) and recombinant bacmid DNA purified. Recombinant baculoviruses expressing XPF and ERCC1 were generated using standard protocols^47^ (Oxford Expression Technologies). In short, 1×10^6^ SF21 cells (Thermo-Fisher) grown in SFIII media (Thermo-Fisher) and 10 μg/ml gentamycin (Life Technologies) were infected at a multiplicity of infection (MOI) of 2 and harvested after 72 h. Cell pellets were resuspended in extract buffer (20 mM HEPES pH 7.8, 150 mM NaCl, 1 mM tris(2-carboxyethyl)phosphine (TCEP), 10% glycerol, 2mM MgCl_2_, 0.01% (w/v) 3-((3-cholamidopropyl) dimethylammonio)-1-propanesulfonate (CHAPS), 0.25 tablet of EDTA-free protease-inhibitor cocktail per litre of culture, and 1 μl per 250 mL lysate BaseMuncher (Expedeon) and lysed by sonication. The lysate was cleared of insoluble cell debris by centrifugation at 35,000g for 45 min and incubated with Strep-tactin resin (GE Healthcare) for 1 h at 4 °C. The resin was extensively washed with extract buffer minus protease inhibitors and BaseMuncher and incubated for 12 h with Tobacco Etch Virus (TEV) protease (in-house). The eluate, containing XPF-ERCC1 was concentrated and further purified using anion-exchange (HiTrap-Q, GE Healthcare) before a final size exclusion chromatography (SEC) step using a Superdex-200i column (GE Healthcare) in cryo buffer (20 mM HEPES pH 7.8, 150 mM NaCl, 1 mM TCEP, 0.01% (w/v) CHAPS). Mutants were cloned using the Q5 site-directed mutagenesis kit (NEB) and were then expressed using the same protocol as described above for wild type XPF-ERCC1.

### XPF-ERCC1 DNA complex assembly

DNA with a modified phosphorothioate backbone (SLp DNA) was resuspended in DNA resuspension buffer (10 mM Tris, pH 7.8, 1 mM EDTA and 75 mM NaCl) and annealed to form a stem-loop structure. Purified XPF-ERCC1 was buffer exchanged into XPF-ERCC1 DNA cryo buffer (20 mM HEPES pH 7.8, 150 mM NaCl, 1 mM TCEP, 0.01% (w/v) CHAPS, 5 mM CaCl_2_, 0.5 mM EDTA) and then incubated with SLp DNA at a 1:2 protein:DNA molar ratio for 10 mins at 4°C followed by crosslinking with 0.05% (v/v) glutaraldehyde for 10 mins at 4 °C. The crosslinking reaction was quenched by the addition of 1 mM Tris-HCl, pH 7.8 and the complex further purified via SEC using a Superdex 200i column. Stem-loop sequence: CAGCG*C*T*U*G*G*TTTTTTTTTTTTTTTTTTTT*C*C*A*A*G*CGCTG, where the asterisk * represents a phosphorothioate backbone.

### XPF-ERCC1 cryo-EM grid preparation and data collection

For cryo-EM analysis, 4 μl of the purified XPF-ERCC1 heterodimer at 1.5 mg/ml was applied to both R1.2/1.3 400 mesh QuantiFoil® grids that had been previously glow-discharged for 45 s at 42 mA. The grids were blotted for 4 s at 100% humidity and 4 °C and plunged into liquid ethane cooled by liquid nitrogen using a FEI Vitrobot MK IV. The grids were then loaded onto a Titan Krios transmission electron microscope operated at 300 kV (Thermo-Fisher). Images were collected in counting mode using a Gatan K2 Summit direct electron detector camera mounted behind a GIF Quantum energy filter operating in zero-loss mode. Exposures were 15 s, with a total dose of 63 e-/Å2 dose-fractionated into 40 frames with a calibrated pixel size of 1.38 Å. Images were recorded with a defocus of 1.5 µm to 4 µm. A total of 15,315 micrographs were collected from three separate data collection sessions.

### XPF-ERCC1 cryo-EM image processing

Movie frames were corrected for motion using MotionCor2^48^, and contrast transfer function was estimated using CTFfind4.1^49^ within Scipion-1.2^50^. The total number of movies collected across three collections was 15,315, of which 14,453 were used. 200 micrographs were selected from the first collection from which 82,412 particles were picked using Xmipp3^51^ semi-automated picking and extracted using RELION-3^52^. The particles were sorted using Xmipp3^51^ screen particles followed by three rounds of reference-free 2D classification in CryoSPARC-2^53^. Asubset of six 2D classes were selected that represented different views of the molecule and used as templates for reference-based particle picking using Gautomatch^54^ on the full dataset. This approach yielded 396,106, 1,201,881 and 2,391,900 particles for data collection runs one, two and three respectively. The particles were extracted and binned two-fold using RELION-3^52^, sorted using Xmipp3^51^ screen particles and then submitted for three rounds of reference-free 2D classification in CryoSPARC-2^53^. This reduced the particle numbers to 151,412, 390,007 and 1,074,111 particles for data collection runs one, two and three respectively. Four initial models were generated using the ab initio reconstruction program in CryoSPARC-2^53^ and were used as references for 3D classification using heterogeneous refinement in CryoSPARC-2^53^. Multiple rounds of heterogeneous refinement yielded 44,312, 126,492 and 390,712 particles in well-defined classes for data collection runs one, two and three respectively. All 561,516 particles from the three collections were re-extracted in an un-binned 200 x 200-pixel box using RELION-3^52^ and merged. The data then underwent 3D classification without alignment in RELION-3^52^ to identify the most stable, high-resolution class. The two classes that displayed the highest-resolution features, comprising 405,339 particles, were refined to 4.1 Å resolution in CryoSPARC-2^53^ using non-uniform refinement. Per-particle motion corrected was carried out using Bayesian polishing in RELION-3^52^. The shiny, polished particles were then refined to 4.0 Å resolution in CryoSPARC-2^53^ using non-uniform refinement.

Inspection of the 4.0 Å resolution map rendered by local resolution in UCSF Chimera^55^ identified the dimeric XPF-ERCC1 2x(HhH)_2_ domain as the lowest resolution region of the map, suggesting some degree of mobility. A mask which excluded the low-resolution XPF-ERCC1 2x(HhH)_2_ was generated in Chimera^55^ and using the particle subtraction tool in CryoSPARC-2^53^ the portion of the particle images aligning to the hairpin density in the map was removed. Non-uniform local refinement in CryoSPARC-2^53^ was performed on the subtracted particles, re-aligning them to the masked reference volume, leading to a reconstruction at 3.6 Å resolution which excluded the hairpin portion of the 4.0 Å map.

All resolutions reported here were determined by Fourier Shell Correlation (at FSC = 0.143) based on the ‘gold-standard’ protocol using a soft mask around the complex density^56^. To avoid over-masking, the masked maps were visually inspected to exclude the possibility of the clipping of electron densities. Additionally, the occurrence of over-masking was monitored by inspecting the shapes of FSC curves. The two half maps had their phases randomised beyond the resolution at which the no-mask FSC drops below the FSC = 0.143 criterion. The tight mask is applied to both half maps, and an FSC is calculated. This FSC is used along with the original FSC before phase randomization to compute the corrected FSC. Local resolution was calculated using Blocres within CryoSPARC-2^53^. For visualisation, maps were sharpened by applying an automated local-resolution weighted negative B factor using the local filtering function of CryoSPARC-2^53^.

### XPF-ERCC1 model building

Initially the crystal structures of the ERCC1 nuclease-like domain (NLD) (PDB code: 2A1I) and the tandem helix-hairpin-helix domains comprising XPF and ERCC1 chains (PDB code: 2A1J) were rigid body fitted into the locally-filtered and sharpened map obtained at 4.0 Å resolution. Homology models were generated for the XPF RecA1 domain using Phyre2 and rigid body fit into the map using the same procedure. Subsequently the fitted domains were rebuilt manually using COOT^57^ optimising the fit where sidechain densities were evident prior to using FlexEM^58^ and real-space refinement as implemented in PHENIX^59^ whilst imposing secondary structural and geometric restraints prevent overfitting. The RecA2 and helical domains were built de novo and subjected to PHENIX^59^ real-space refinement. A further 6 cycles of re-building and refinement in COOT^57^ and PHENIX^59^ lead to a model containing 743 residues from XPF and 195 from ERCC1. Linkers regions connecting the XPF nuclease and ERCC1 NLD domains to their respective (HhH)_2_ domains were built manually into the map and the N-terminal portion of the XPF nuclease domain homology model was rebuilt in COOT^57^ to fit the map. The final atomic model was evaluated using MolProbity^60^.

### XPF-ERCC1-DNA complex cryo-EM grid preparation and data collection

XPF-ERCC1-DNA complex was concentrated to 1.3 mg/ml and applied to Quantifoil R1.2/1.3 300 mesh copper grids. The freezing and imaging conditions used were the same as for the DNA-free XPF-ERCC1 complex described above and 8965 movies were collected.

### XPF-ERCC1-DNA complex cryo-EM image processing

Motion correction and CTF estimation was performed as previously described for the XPF-ERCC1 data collections. 7982 micrographs were manually selected for processing. Particle picking was carried out as described for the XPF-ERCC1 data collections. 3,432,565 particles were extracted and sorted using Xmipp3^51^ screen particles and then submitted for six rounds of reference-free 2D classification in CryoSPARC-2^53^. 688,821 particles were used to generate 4 ab initio reconstructions which were then used as references for 3D classification using heterogeneous refinement in CryoSPARC-2^53^. Multiple rounds of heterogeneous refinement were carried out yielding one well-ordered reconstruction comprising 199,022 particle images. This class was refined to 7.7 Å resolution using non-uniform refinement in CryoSPARC-2^53^. A mask was generated using UCSF Chimera^55^ that excluded both the DNA and hairpin domain density which was used to carry out masked refinement improving the resolution of the sub-volume to 5.9 Å.

### XPF-ERCC1-DNA complex model building

Individual domains of XPF-ERCC1 were taken from the DNA-free structure and fitted into the DNA-bound cryo-EM map density as rigid bodies using the UCSF Chimera^55^ fit-in-map tool. The nuclease domain dimer of the A. pernix XPF homolog (PDB:2BGW) bound to dsDNA was fitted into the DNA-bound map density and the subsequent position of the DNA-bound A. pernix hairpins used as a reference to align the human hairpin domain using MatchMaker in UCSF Chimera^55^. The DNA from the A. pernix structure was reduced to a 10 base-pair duplex and modelled into the map whilst preserving the hairpin domain-DNA contacts. The sequence conservation of the functional human ERCC1 and A. pernix (HhH)_2_ domains is high: 25.5% identical and 69.1% similar residues. The ds-RNA bound structure of MDA5 was placed into the DNA-bound map density as a guide to place the helical domain of XPF by inspecting the position of the homologous domain in MDA5.

### XPF-ERCC1-DNA-TFIIH-XPA complex modelling

The XPF-ERCC1-DNA structure was aligned to the TFIIH-DNA-XPA structure (PDB code: 6RO4) through structural super-imposition in UCSF Chimera^55^ and alignment with the two DNA strands of a single duplex from each structure. The ds/ss DNA junction was defined by the high-resolution DNA structure in the TFIIH-XPA complex and demarcated by the position of the XPA β-hairpin. The structure was subsequently manually refined in Coot^57^.

### XPF ERCC1 cross-linking mass spectrometry

All chemicals were purchased from Sigma-Aldrich unless otherwise stated. 100 µg XPF-ERCC1 heterodimer at a concentration of 1 mg/ml in 20 mM HEPES, pH 7.8, 10% Glycerol (w/v), 0.01% CHAPS (w/v), 150 mM NaCl, 1 mM TCEP, 0.5 mM EDTA was cross-linked using 1 mM disuccinimidyl sulfoxide (DSSO) (Thermo-Fisher) with mild shaking for 30 min at 37 ^o^C. The reaction was quenched using a final concentration of 50 mM ammonium bicarbonate for a further 20 mins at 37 °C. To remove potential aggregates, gradient ultra-centrifugation was employed using a 5-30% Glycerol gradient in 20 mM Hepes, 150 mM NaCl, mixed using a Gradient Master (BioComp), and centrifuged for 16 h at 4 ^o^C at 40,000 RPM using a SW 55 Ti Rotor (Beckman Coulter)^61^. 100 µL fractions were collected and silver stained to identify fractions containing cross-linked non-aggregated XPF-ERCC1. Fractions containing cross-linked proteins were then pooled and buffer exchanged into 8 M urea using a Vivaspin 500, 30,000 molecular weight cut off (MWCO) PES filter (Sartorius, VS0122). Cysteine reduction was carried out using 2.5 mM TCEP for 30 min at 37 ^o^C and alkylated in the dark using 5 mM iodoacetamide at room temperature. The urea was then buffer exchanged for 50 mM ammonium bicarbonate and proteins were proteolysed using trypsin (Promega) at 1:50 (w/w) trypsin:protein overnight at 37 ^o^C. The solution was acidified using 2 % formic acid and peptides were the spun through the MWCO filter and desalted using in-house built STAGE tips made using Empore SPE C18 disks (3M, 66883-U). The eluent was then dried to completion. Peptides were reconstituted in 0.1% trifluoroacetic acid (TFA) and chromatographically resolved using an Ultimate 3000 RSLCnano (Dionex) HPLC. Peptides were first loaded onto an Acclaim PepMap 100 C18, 3 µm particle size, 100 Å pore size, 20 mm x 75 µm ID (Thermo Scientific, 164535) trap column using a loading buffer (2 % acetonitrile (MeCN) and 0.05 % TFA) with a flow rate of 7 µL/minute. Chromatographic separation was achieved using an EASY-Spray column, PepMap C18, 2 µm particles, 100 Å pore size, 500 mm x 75 µm ID (Thermo Scientific, ES803). The gradient utilised a flow of 0.3 µl/minute, starting at 98 % mobile A (0.1 % formic acid, 5 % dimethyl sulfoxide (DMSO) in H_2_O) and 2 % mobile B (0.1 % formic acid, 75 % MeCN, 5 % DMSO and 19.9 % H_2_O). After 6 min, mobile B was increased to 30 % over 69 min, to 45 % over 30 min, further increased to 90 % in 16 min and held for 4 min. Finally, Mobile B is reduced back to 5 % over 1 min for the rest of the acquisition. Data was acquired in real time over 140 minutes using an Orbitrap Fusion Lumos Tribrid mass spectrometer in positive, top speed mode with a cycle time of 5 s. The chromatogram (MS1) was captured using 60,000 resolution, a scan range of 375-1500 with a 50 ms maximum injection time, and 4e5 AGC target. Dynamic exclusion with repeat count 2, exclusion duration of 30 s, 20 ppm tolerance window was used, along with isotope exclusion, a minimum intensity exclusion of 2e4, charge state inclusion of 3-8 ions and peptide mono isotopic precursor selection. Precursors within a 1.6 m/z isolation window were then fragmented using 25% normalised CID, 100 ms maximum injection time and 5e4 AGC target. Scans were recorded using 30,000 resolution in centroid mode, with a scan range of 120 – 2000 m/z. Spectra containing peaks with a mass difference of 31.9721 Da were further fragmented with a 30% normalised higher collision induced dissociation, using a 2 m/z isolation window, 150 ms maximum injection time and 2e4 AGC target. Four scans were recorded using an ion trap detection in rapid mode starting at 120 m/z.

### XL-MS data analysis

Data processing was carried out using Proteome Discoverer Version 2.2 (Thermo Scientific) with the XlinkX^62^ node. The acquisition strategy was set to MS2_MS3 mode. The database comprised solely of the specific XPF and ERCC1 sequences. Trypsin was selected as the proteolytic enzyme allowing up to two missed cleavages with a minimal peptide length of five residues. Masses considered were in the range of 300-10000 Da. The precursor mass tolerance, FTMS fragment mass tolerance, and ITMS Fragment Mass Tolerance were set to 10 ppm, 20 ppm and 0.02 Da respectively. A static carbamidomethyl (+57.021 Da) modification was utilised for cysteine residues, with additional dynamic modifications considered including; amidated and hydrolysed DSSO (+142.050 Da and +176.014 Da, respectively) on lysine serine and threonine residues, oxidation (+15.995 Da) on methionine residues, and protein N-terminal acetylation (+42.011 Da). The FDR threshold was set to one with the strategy set to simple. The list of reported cross-linked spectral matches were manually examined and cross-links with spectra that did not contain acceptable b and y ion coverage were excluded.

### XPF-ERCC1-SLX4^NTD^ Complex Assembly

cDNA encoding the SLX4^NTD^ (residues 1-758) (NCBI reference sequence: NM_032444) was shuttled into a pGEX-1 vector (Sigma). Recombinant baculoviruses expressing the SLX4^NTD^ were generated as previously described and used to infect 1×10^6^ SF21 cells (Thermo-Fisher) grown in SFIII media (Thermo-Fisher) and 10 μg/ml gentamycin (Life Technologies) at an MOI of 0.5. These cells were co-infected with XPF-ERCC1 expressing baculovirus at an MOI of 2. Cells were pelleted after 72 h and protein extracted as previously described for XPF-ERCC1. Following strep-tactin affinity purification, the complex was purified using anion-exchange (HiTrap-Q, GE Healthcare) using a gradient of 150 mM NaCl to 500 mM NaCl over 20 ml of extract buffer minus protease inhibitors and Base-Muncher. This separated the SLX4^NTD^-XPF-ERCC1 complex from unbound XPF-ERCC1. Fractions containing the SLX4^NTD^-XPF-ERCC1 complex were pooled and concentrated prior to a final size exclusion chromatography (SEC) step using a Superose-6 increase column (GE Healthcare). Fractions containing both XPF and SLX4^NTD^ were identified via Western blot.

### Real-time fluorescence incision assay

Fluorescently labelled stem-loop (SLF) DNA substrates, containing a 5′ 6-FAM fluorophore and 3′-BHQ1 quench, were purified by SEC (Superdex-200i, GE Healthcare) in assay buffer (5 mM HEPES, 10 % glycerol, 0.5 mM DTT, 1 mM MnCl_2_ and 40 mM NaCl. The purified substrates were then annealed by heating to 95 °C for 1 min followed by cooling to 4 °C and dispensed into the assay plate. Reactions were carried out in 384-well black, flat-bottomed microtitre plates (Corning 3854). Purified XPF-ERCC1 was buffer exchanged into assay buffer and 5 nM added to each in a total volume of 20 µl to initiate the endonuclease reaction. Fluorescence measurements were carried out using the CLARIOstar plate reader (BMG Labtech) using an excitation wavelength of 483 nm and an emission wavelength of 525 nm. 60 readings were collected at 30 s intervals and the linear response range for each substrate was used to determine the change in fluorescence per unit time. Kinetic parameters were calculated using the Michaelis–Menten equation in PRISM7. Experimental product release was quantified by plotting the relative fluorescence units (RFU) produced by known amounts of the cleavage products against their concentration to generate a standard curve. SLF sequence: 6-FAM-5’-CAGCGCTUGGTTTTTTTTTTTTTTTTTTTTCCAAGCGCTG-3’-BHQ1 Cleavage product #1: 6-FAM-5’-CAGCGCTC 3’ Cleavage product#2:GGTTTTTTTTTTTTTTTTTTTTC-CGAGCGCTG-3′-BHQ1

## Data Availability

The coordinates for the DNA-free XPF-ERCC1 structure are available in the PDB with the primary accession code 6SXA and EMDB entry code EMD-10337. The coordinates for the DNA-bound XPF-ERCC1 structure are available in the PDB with the primary accession code 6SXB and EMDB entry code EMD-10338.

## Acknowledgments

We thank members of the McDonald laboratory for helpful discussions and comments on the manuscript, in particular Amy Whittaker who also assisted in running the activity assays. We thank Andrew Purkiss of the structural biology science technology platform for assistance in refining the structures and Raffaella Carzaniga for EM training and support. We also thank Steve West for an anti-SLX4 antibody and Rick D Wood (MD Anderson Centre, Texas) for stimulating and encouraging the project in its early phases. We acknowledge the helpful advice and support of Peter Rosenthal on all aspects of cryo-electron microscopy. M.J. was funded by a Crick/UCL joint PhD studentship. N.Q.M. acknowledges that this work was supported by the Francis Crick Institute, which receives its core funding from Cancer Research UK (FC001115), the UK Medical Research Council (FC001115) and the Wellcome Trust (FC001115). EPM and FB acknowledge support from Cancer Research UK (C12209/A16749).

## Author Contributions

M.J. cloned and purified XPF-ERCC1 complexes, processed all cryo-EM data, carried out activity assays and built the atomic models. M.J. and F.B. carried out EM grid preparations. F.B. and C.E assisted in grid screening, A.N and P.R. ran the data collections. F.B. and E.P.M. assisted the XPF-ERCC1 single particle reconstruction and interpretation. D.B. assisted in model refinement and validation. M.B. for supporting the project in its early phases. A.B. carried out the XL-MS. M.J. and N.Q.M. designed the study, interpreted the results and wrote the paper.

