## Supplementary Data for "Cryo-EM Structures of the XPF-ERCC1 Endonuclease Reveal an Auto-Inhibited Conformation and the Basis for Activation"

### SUPPLEMENTARY FIGURES, LEGENDS AND TABLES

### SUPPLEMENTARY FIGURES

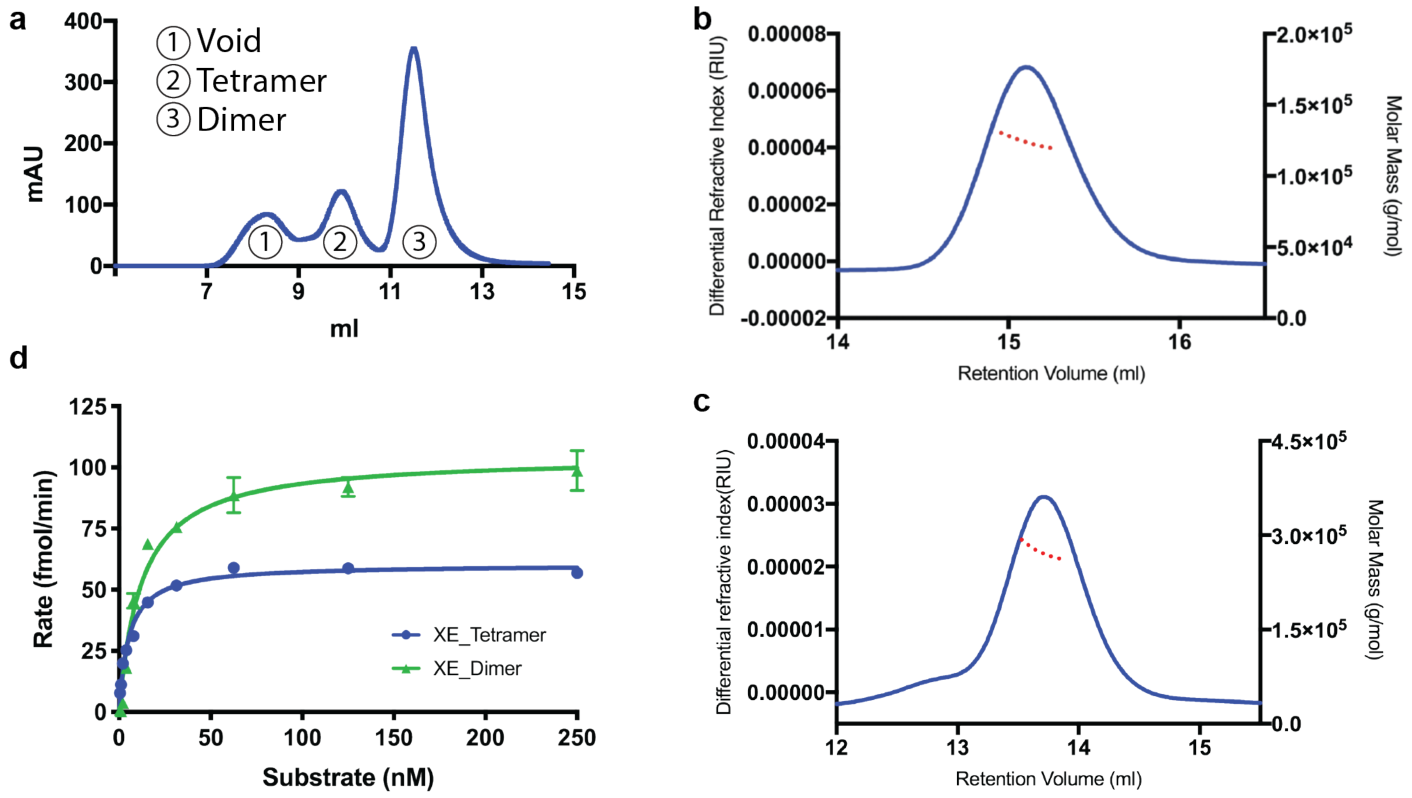


**Figure S1. XPF-ERCC1 purification characterisation.** Recombinant full-length human XPF-ERCC1 heterodimer was produced using the baculovirus expression system in SF21 insect cells[63]. The sample was purified to homogeneity using a twin-Strep affinity-tag, anion exchange and size-exclusion chromatography. **a** Superdex-200 increase (SD200i) column trace for the XPF-ERCC1 complex following affinity and anion exchange purification. Peak 1 at 7.5 ml is aggregated protein, peak 2 at 10 ml is heterotetrameric XPF-ERCC1, peak 3 at 11.5 ml is heterodimeric XPF-ERCC1. Samples from the heterodimer fraction were used for structural studies. **b-c** SEC-MALLS Superose-6 column traces with the estimated molecular mass overlaid in red. **b** XPF-ERCC1 heterodimer with molecular mass of 128 kDa, approximately 9 kDa less than the full-length molecular weight expected most likely due to the proteolytic cleavage of the amino-terminus of ERCC1 (~ 1-95 amino acids). **c** XPF-ERCC1 heterotetramer with molecular mass of 290 kDa. **d** Michaelis-Menten kinetics plot of rate vs stem-loop substrate concentration for the XPF-ERCC1 heterodimer (green) and heterotetramer (blue). Heterodimer Vmax – 104.6 fmol/min, Km – 24.2 fmol, kcat – 10.5 min-1 . Heterotetramer kmax – 60.1 fmol/min, Km – 21.0 fmol, kcat -12.1 min-1.

a

b

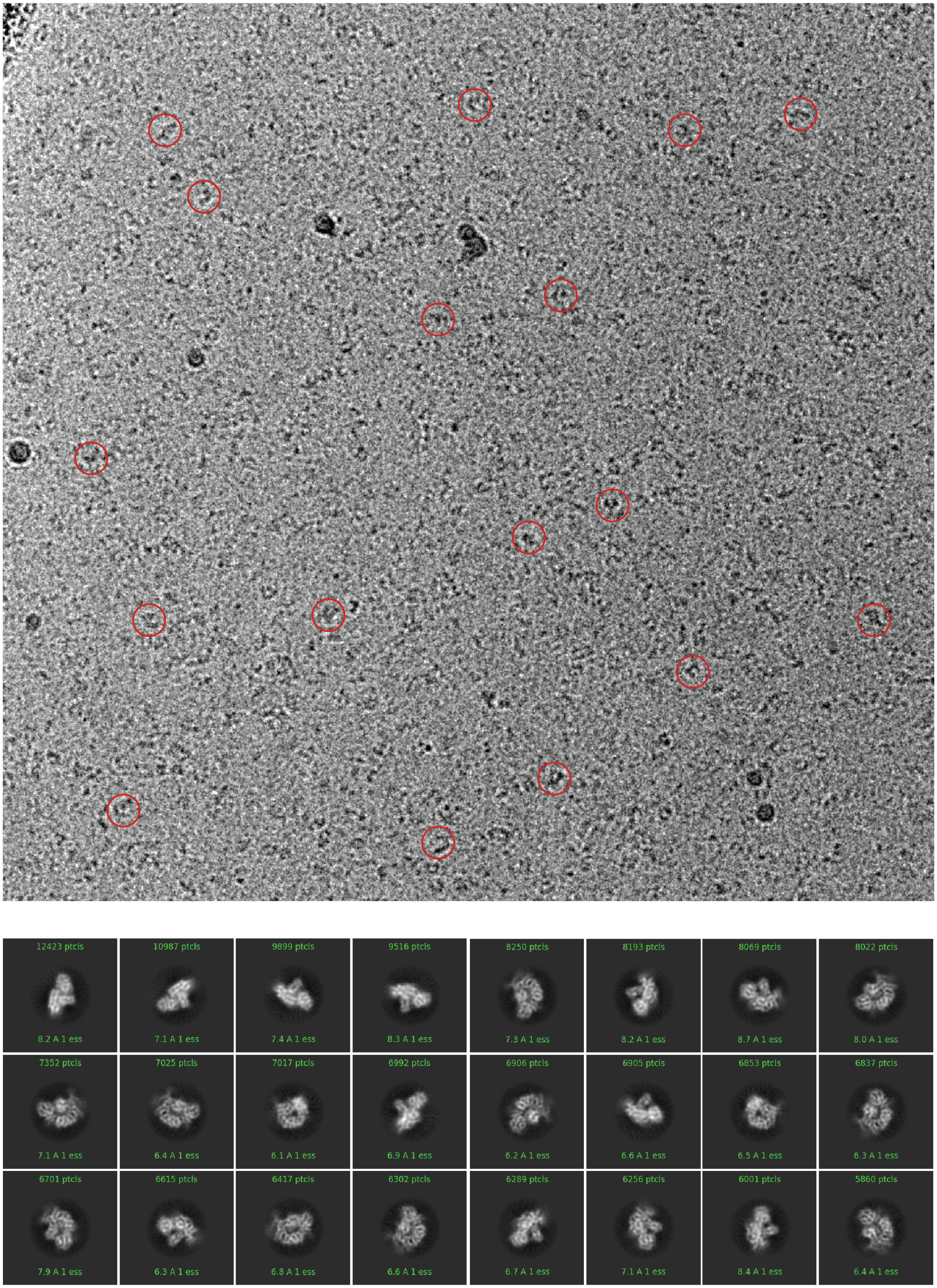


**Figure S2. XPF-ERCC1 cryo-EM micrograph and 2D class averages. a** Motion-corrected cryo-EM micrograph of the XPF-ERCC1 heterodimer. Representative selection of particle images picked in red circles with diameter 160 Å. **b** Representative selection of cryo-EM 2D class-averages of XPF-ERCC1 calculated using CRYOSPARC-2[53] after final 3D classification. A mask with a diameter of 140 Å was applied.

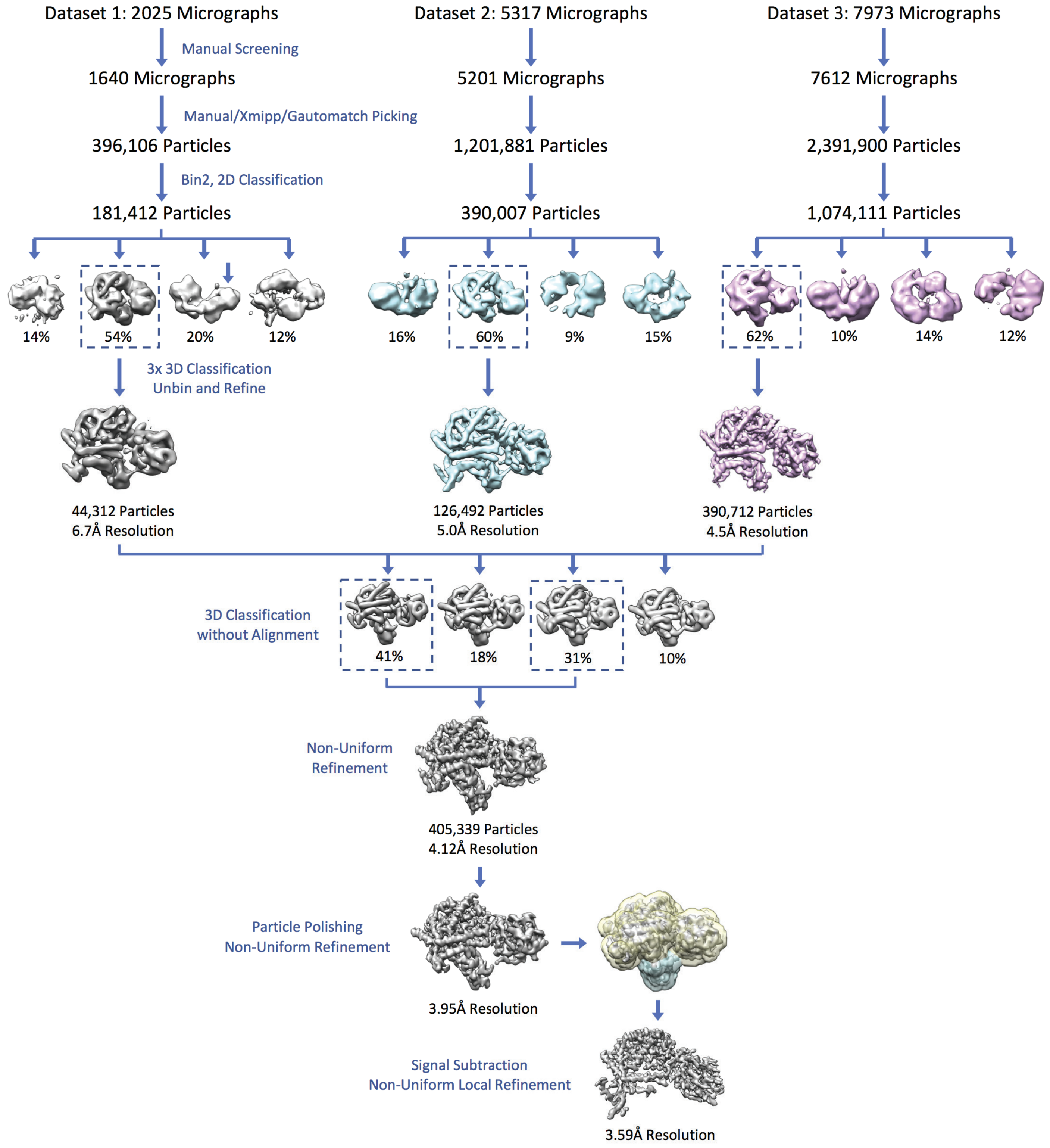


**Figure S3. XPF-ERCC1 cryo-EM processing workflow. a** Datasets one, two and three were initially processed independently. Micrographs were manually screened for crystalline ice and then particle images extracted following picking. After multiple rounds of 2D and 3D classification in CRYOSPARC-2[53] the data were merged and subjected to 3D classification without alignment in RELION-3[52]. Particles belonging to the two highest resolution classes were selected and refined followed by polishing in RELION-3[52]. The data were refined to 3.95 Å in CRYOSPARC-2[53]. Density corresponding to the dimeric hairpin domain was subtracted from the particle images and the remaining density was locally refined to 3.6 Å resolution.

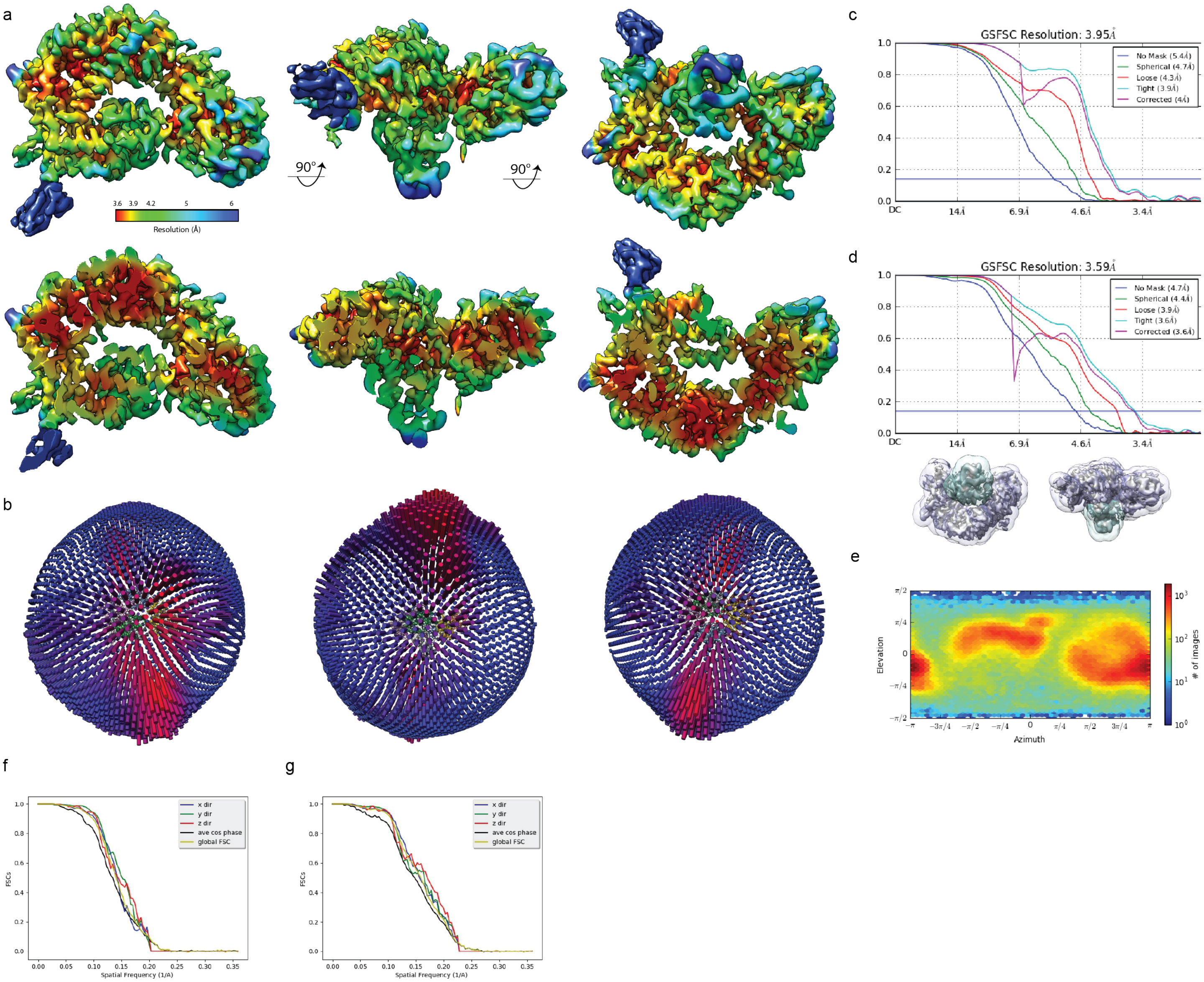


**Figure S4. XPF-ERCC1 local resolution and reconstruction quality. a** XPF-ERCC1 composite cryo-EM map where XPF RecA1, RecA2, helical and nuclease domains and ERCC1 NLD are derived from local refinement without the $2x(HhH)_2$ domain calculated at 3.6 Å, whereas the $2x(HhH)_2$ domain was obtained from global refinement at 4.0 Å resolution. Top, three views of cryo-EM map density filtered by local resolution. Bottom, same views of a slice through map density. **b** Distribution of particle images contributing to global reconstruction in RELION-2[64] displayed in 3D. **c** Fourier-shell correlation (FSC) curve for global refinement at 4.0 Å resolution. FSC = 0.5. **d** FSC curve for local refinement at 3.6 Å resolution. FSC = 0.5. Dip in FSC curves c and d at 6.9 Å is an artefact of the boundary between where randomised phases and where they are experimental. **e** Distribution of particle images contributing to global reconstruction in CRYO-

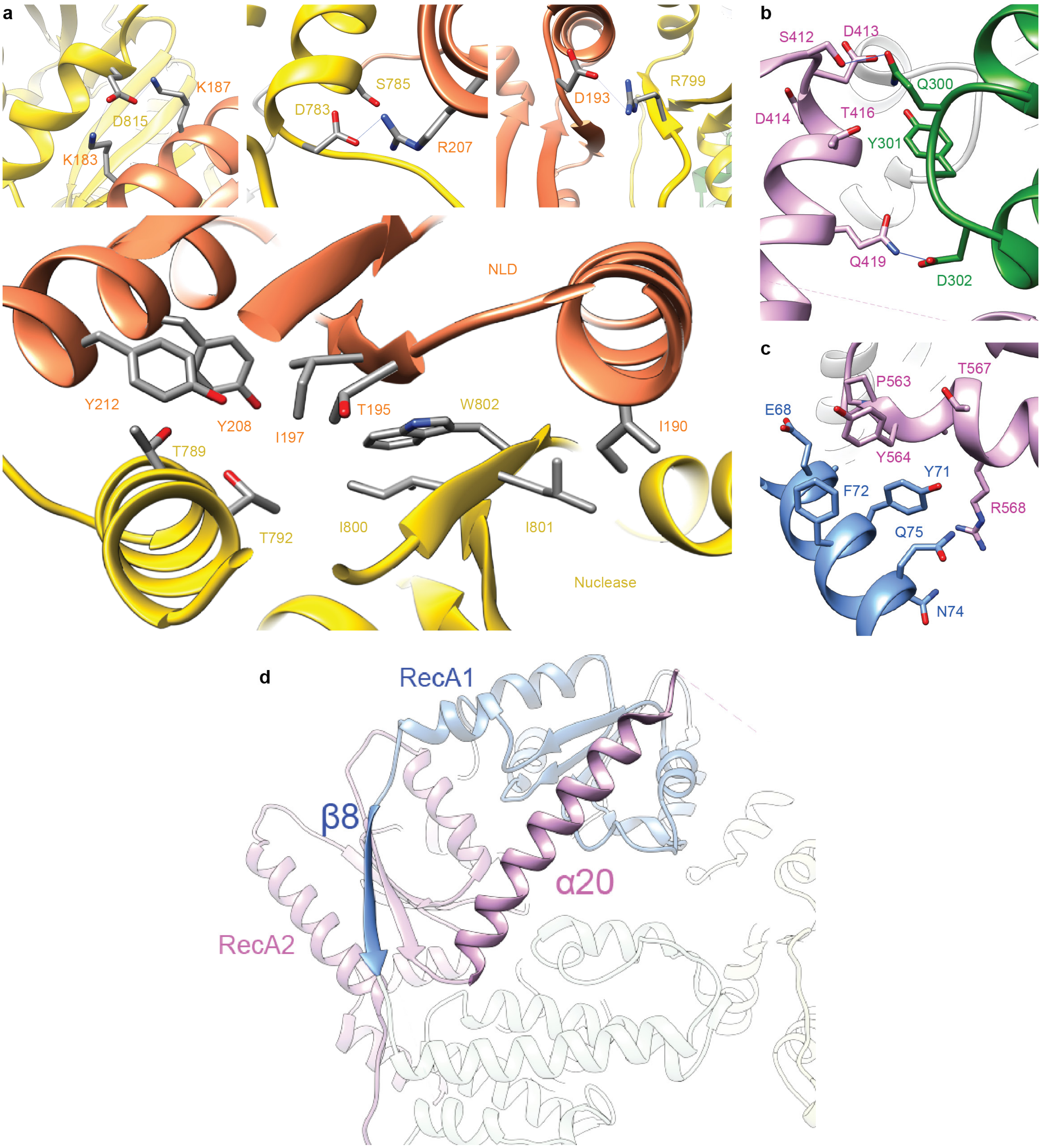


**Figure S5. Additional inter-domain contacts.** Residues displayed as sticks and coloured by heteroatom, grey – C, blue – N, red – O. Hydrogen bonds indicated by blue lines between residues. **a** XPF nuclease (gold) – ERCC1 NLD (coral) interface. Top three images show the salt bridges formed at the periphery of the interface. Bottom, hydrophobic residues at the core of the interface. **b** RecA2 (purple) – helical domain (green) interface highlighting the hydrogen bonds formed between S412 and Q300 and Q419 and D302. **c** RecA1 (blue) – RecA2 (purple) interface highlighting ring stacking between Y71 and Y654. **d** RecA1/A2 domain dimer. Key intertwined secondary structure elements highlighted.

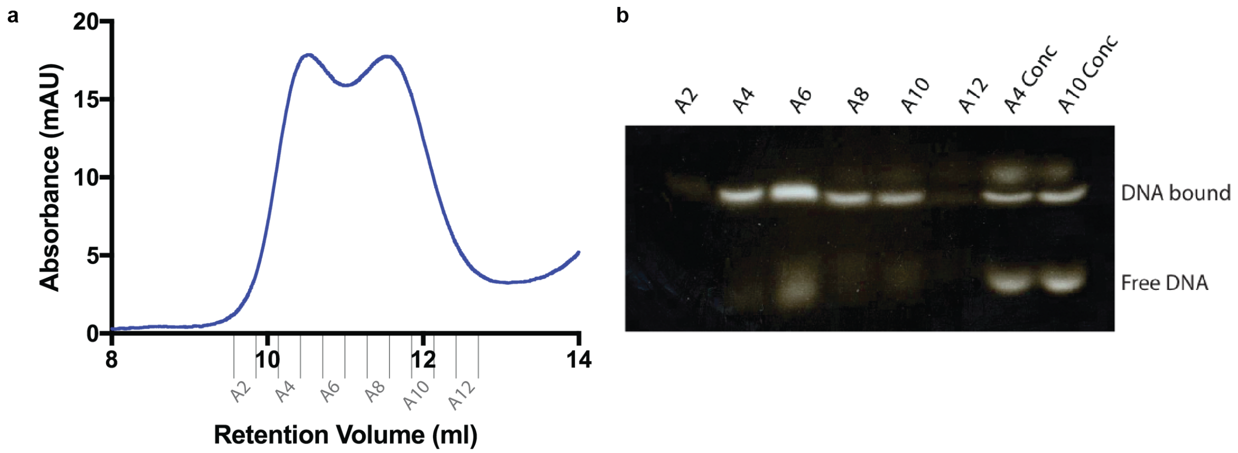


**Figure S6. XPF-ERCC1 DNA binding. a** SD200i SEC trace of XPF-ERCC1-DNA complex. mAU (A280 nm) in blue on the Y axis. Fractions indicated in grey on the X axis. **b** Electrophoretic mobility shift assay (EMSA) showing XPF-ERCC1 mass shift upon binding stem-loop DNA. Fractions A2-A12 from a SD200i SEC trace were assayed for DNA binding following mild crosslinking with BS3. Samples A4 and A10 were concentrated to 1.5 mg/ml and run on the final two lanes. DNA-bound complex and free XPF-ERCC1 indicated by text. Fraction A10 was used for cryo-EM analysis.

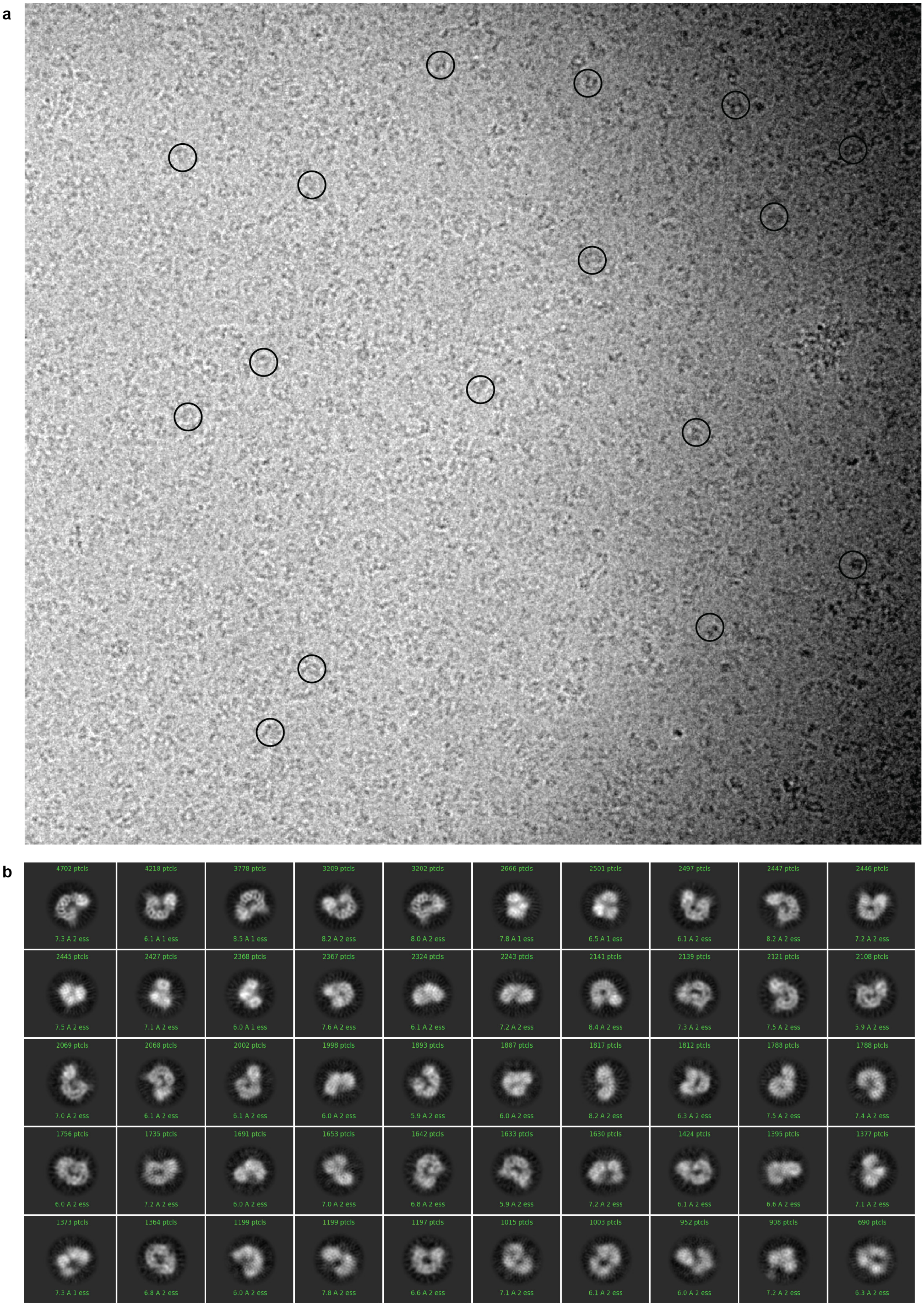


**Figure S7. XPF-ERCC1-DNA cryo EM micrograph and 2D class averages. a** Motion corrected cryo-EM micrograph of the XPF-ERCC1-DNA complex acquired with a Titan Krios microscope at 300 keV, 1.38 Å pixel size and 3.4 µm defocus. Representative particles picked in red circles diameter 150 Å **b** Representative selection of cryo-EM 2D class-averages of XPF-ERCC1-DNA. 199,022 particle images calculated using CRYOSPARC-2[53], 135 Å mask.

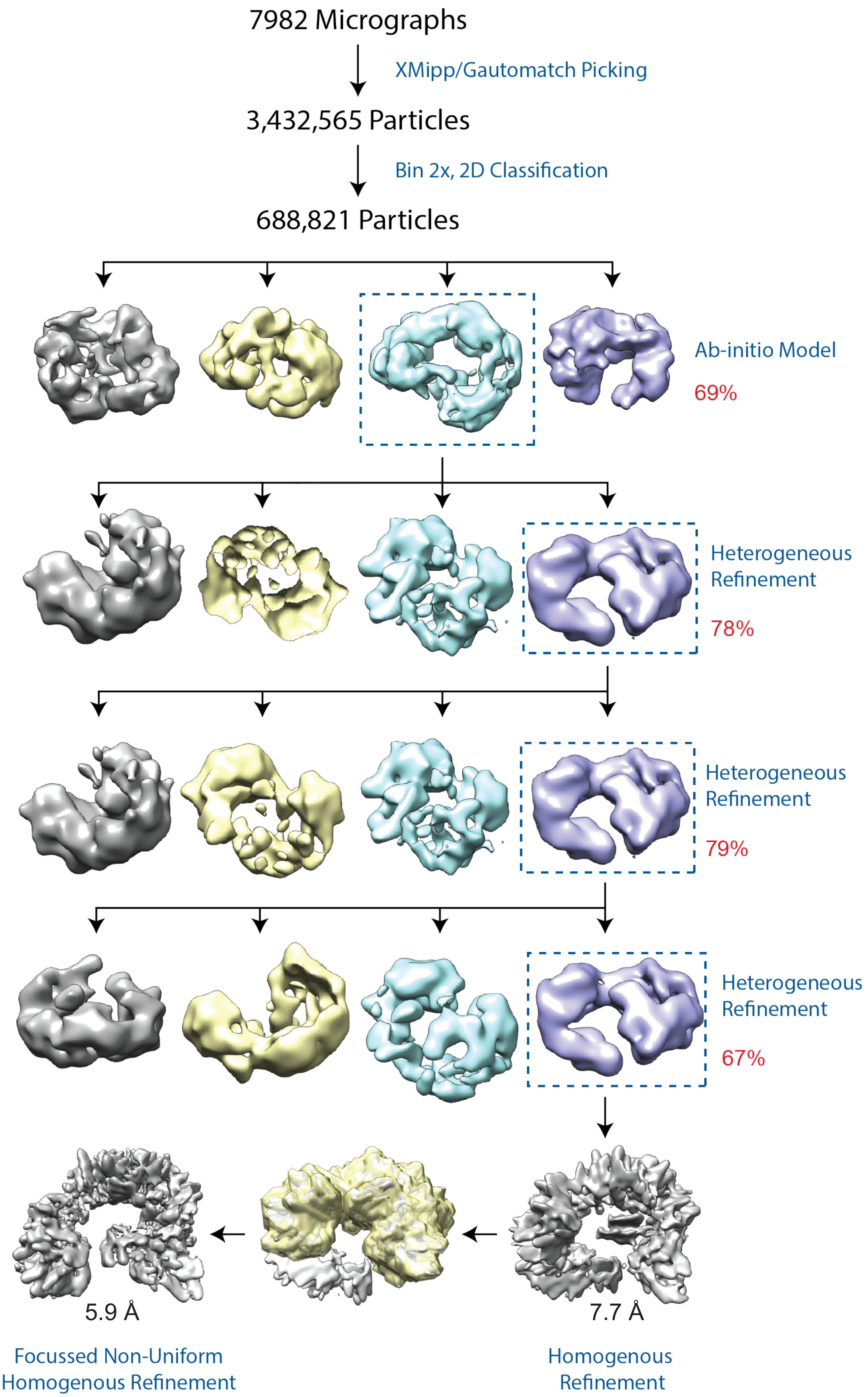


**Figure S8. XPF-ERCC1-DNA cryo-EM processing workflow.** Micrographs were manually screened for crystalline ice and then particle images extracted following picking. After 5 rounds of 2D classification the particle images displaying high-resolution features were classified in 3D using heterogeneous refinement in CRYOSPARC-2[53] using a 3D reference generated by ab-initio reconstruction. The 3D class with the correct volume was selected after 3 rounds of 3D classification and refined to 7.7 Å in CRYOSPARC-2[53]. Density corresponding to the dimeric hairpin domain was subtracted from the particle images and the remaining density was locally refined to 5.9 Å resolution.

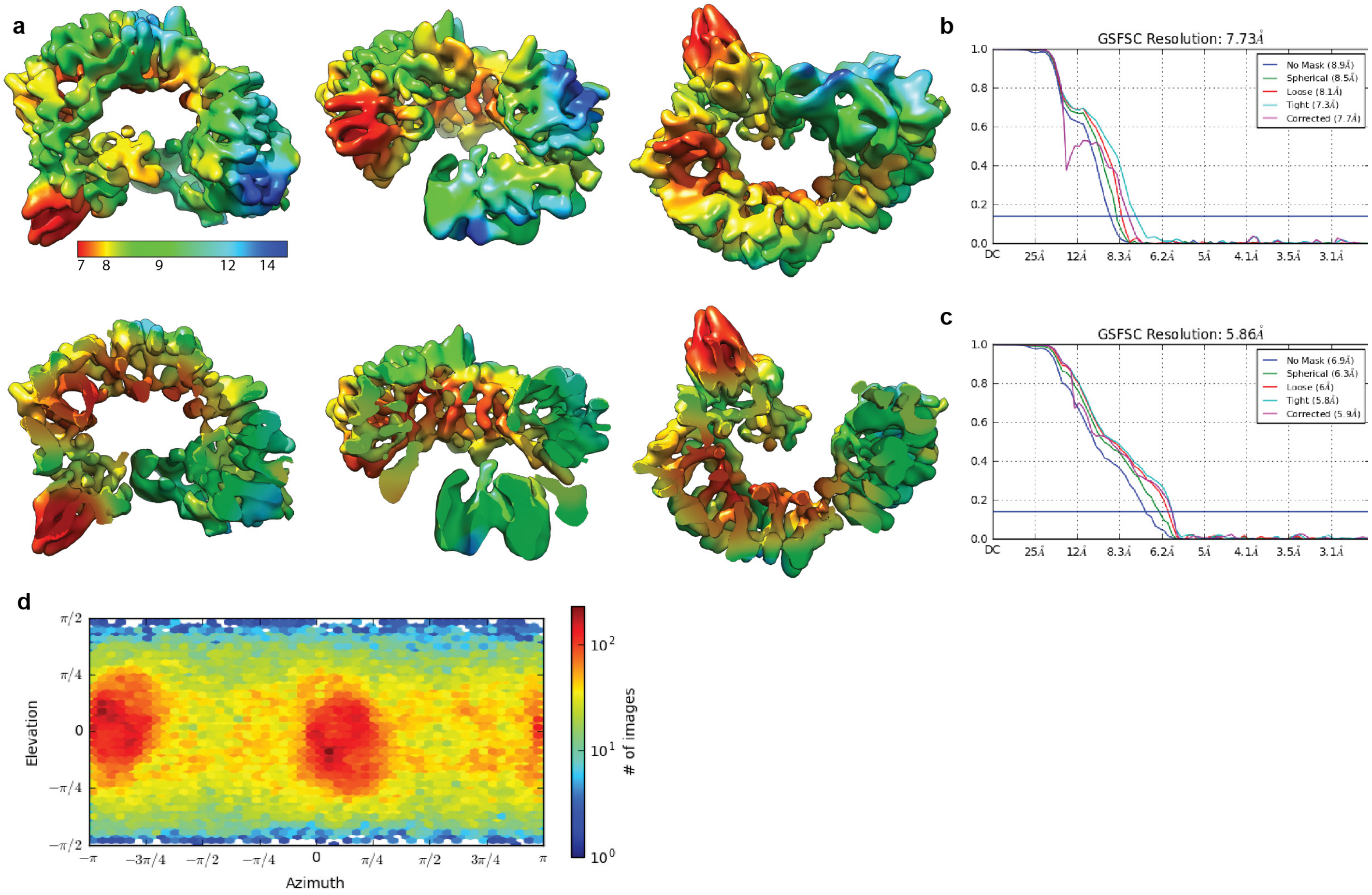


**Figure S9. XPF-ERCC1-DNA local resolution and reconstruction quality. a (Top)** Three orthogonal views of density of the XPF-ERCC1-DNA composite map rendered based on local resolution. **(Bottom)** same three orthogonal views but with density cut-away to reveal inner map features. XPF RecA1, RecA2, helical and nuclease domains and ERCC1 NLD derived from local refinement minus hairpins and DNA at 5.9 Å. XPF and ERCC1 hairpins and DNA from global refinement at 7.7 Å resolution. **b** Fourier-shell correlation (FSC) curve for global refinement at 7.7 Å resolution. **c** FSC curve for local refinement at 5.9 Å resolution. **d** Distribution of particle images contributing to global reconstruction in CRYOSPARC-2[53] displayed in 2D.

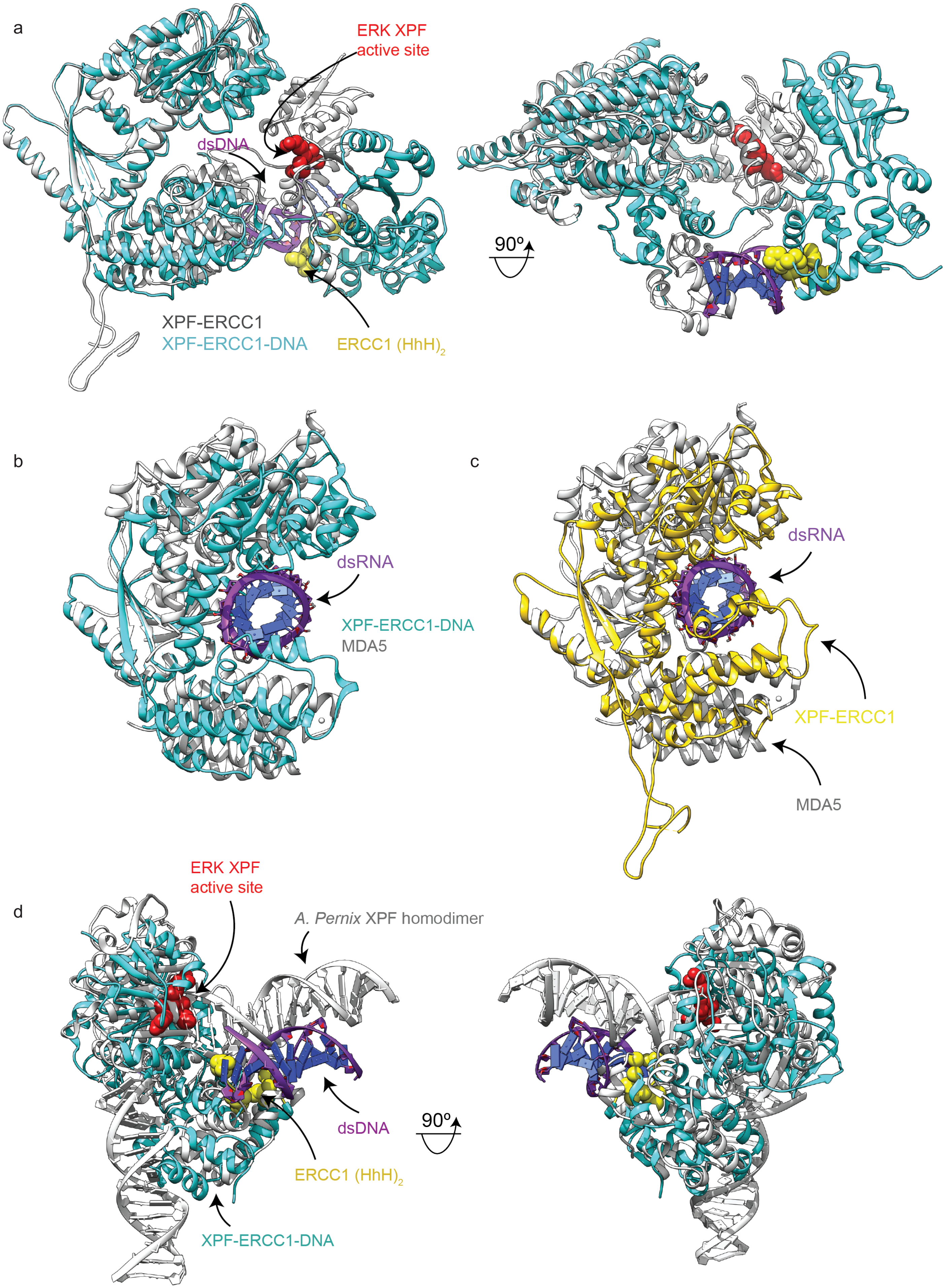
a
ERK XPF
active site
dsDNA
XPF-ERCC1
XPF-ERCC1-DNA
ERCC1 (HhH)2
90º
b
dsRNA
XPF-ERCC1-DNA
MDA5
c
dsRNA
XPF-ERCC1
MDA5
d
ERK XPF
active site
A. Pernix XPF homodimer
dsDNA
ERCC1 (HhH)2
XPF-ERCC1-DNA
90º

**Figure S10. Structural comparisons of XPF-ERCC1 with homologues. a** Structural comparison of XPF-ERCC1 in the presence and absence of stem-loop DNA. DNA-free XPF-ERCC1 in grey, overlaid with stem-loop DNA-bound XPF-ERCC1 in cyan. Labels indicate location of key features on the XPF-ERCC1-DNA model. **b** Structural comparison of the HLM of the XPF-ERCC1-DNA complex with dsRNA-bound MDA5 (PDB code:4GL2). MDA5 was manually aligned to the XPF RecA2 domain beta sheet. DNA-free XPF-ERCC1 in cyan overlaid with dsRNAbound MDA5 in grey. dsRNA from MDA5 in purple. **c** Structural comparison of the HLM of the DNA-free XPF-ERCC1 complex with dsRNA-bound MDA5 (PDB code:4GL2). DNA-free XPF-ERCC1 model in yellow, dsRNA-bound MDA5 in grey. dsRNA from MDA5 in purple. α17 from DNA-free XPF-ERCC1 in green. By superimposing the RecA2 sheet between XPF (residues 406-425) and MDA5 (PDB code: 4GL2, residues 720-739) the RecA1 domain is rotated by 25° relative to the equivalent RecA domain from the dsRNA-bound MDA5 (rmsd 11.9Å). The RecA1 ß-sheet is translated by ~15Å from the MDA5 equivalent RecA1 ß-sheet. The RecA2 domain α17 helix orientation is perpendicular to the direction of the equivalent helix (residues 742-746) of MDA5. The positioning of α17 locks the XPF helical domain in place, rotated by 45° relative to the equivalent domain in MDA5, in a conformation that is not conducive to dsDNA-binding. **d** Structural comparison of dsDNA-bound A. pernix XPF homodimer with the human XPF-ERCC1-DNA complex. XPF-ERCC1-DNA in cyan, A. pernix XPF in grey. Two structures aligned by their 2x$(HhH)_2$ domains. Labels indicate location of key features on the XPFERCC1-DNA model.

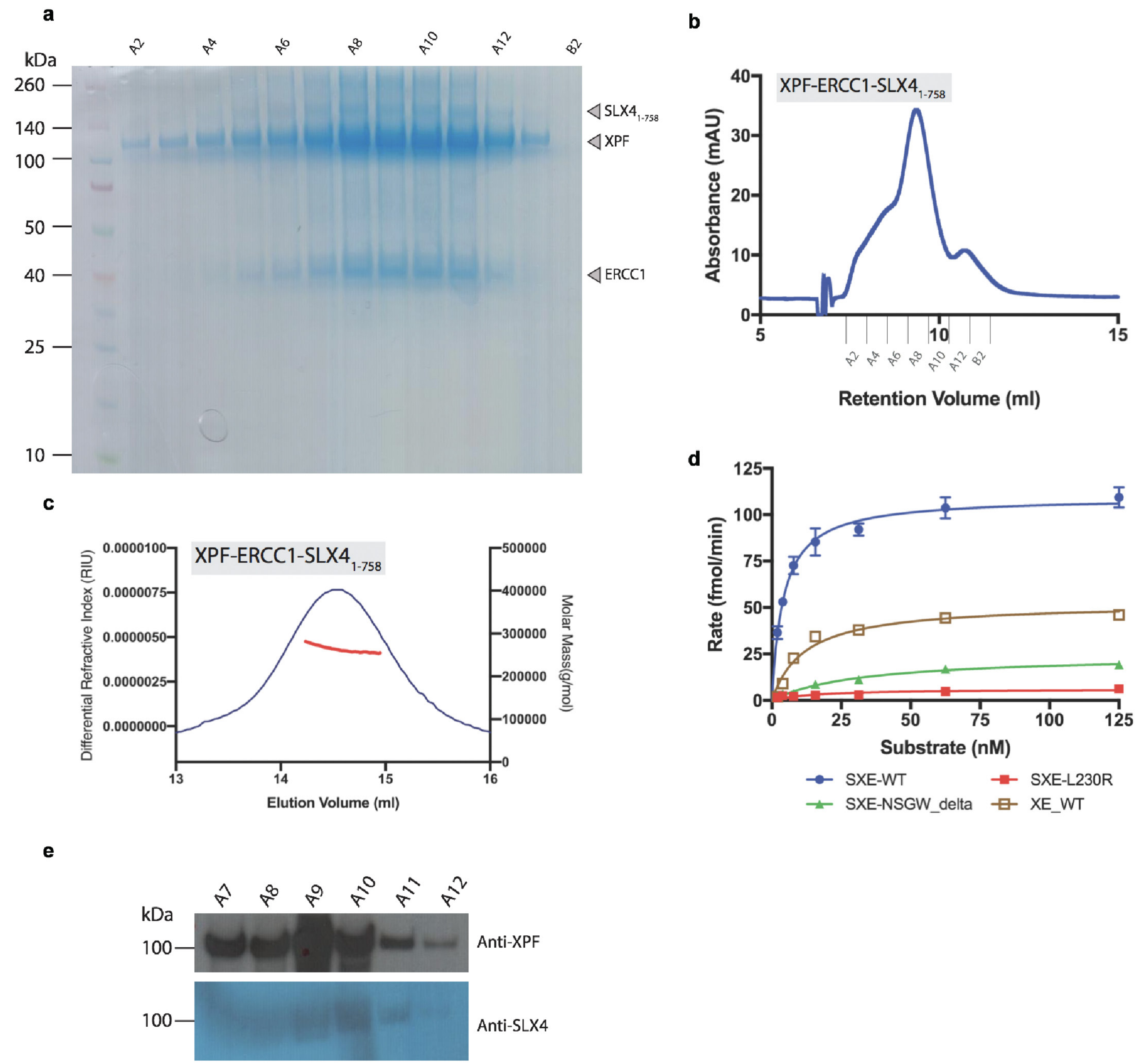


**Figure S11. Purification and characterisation of XPF-ERCC1-SLX4$^{NTD}$ activity and stoichiometry. a** SDS-PAGE gel of the XPF-ERCC1-SLX4$^{NTD}$ complex. **b** Superose-6 increase SEC column trace for the XPF-ERCC1-SLX4NTD complex. **c** SEC-MALLS for the XPF-ERCC1-SLX4$^{NTD}$ complex. Molecular weight 240 kDa, indicating a 1:1:1 complex. **d** Michealis-Menten plot of rate vs stem-loop substrate concentration for XPF-ERCC1 wild-type XPF-ERCC1. Kinetic values in Table 3. **e** Western blot showing fractions with both ERCC1 and SLX4 present.

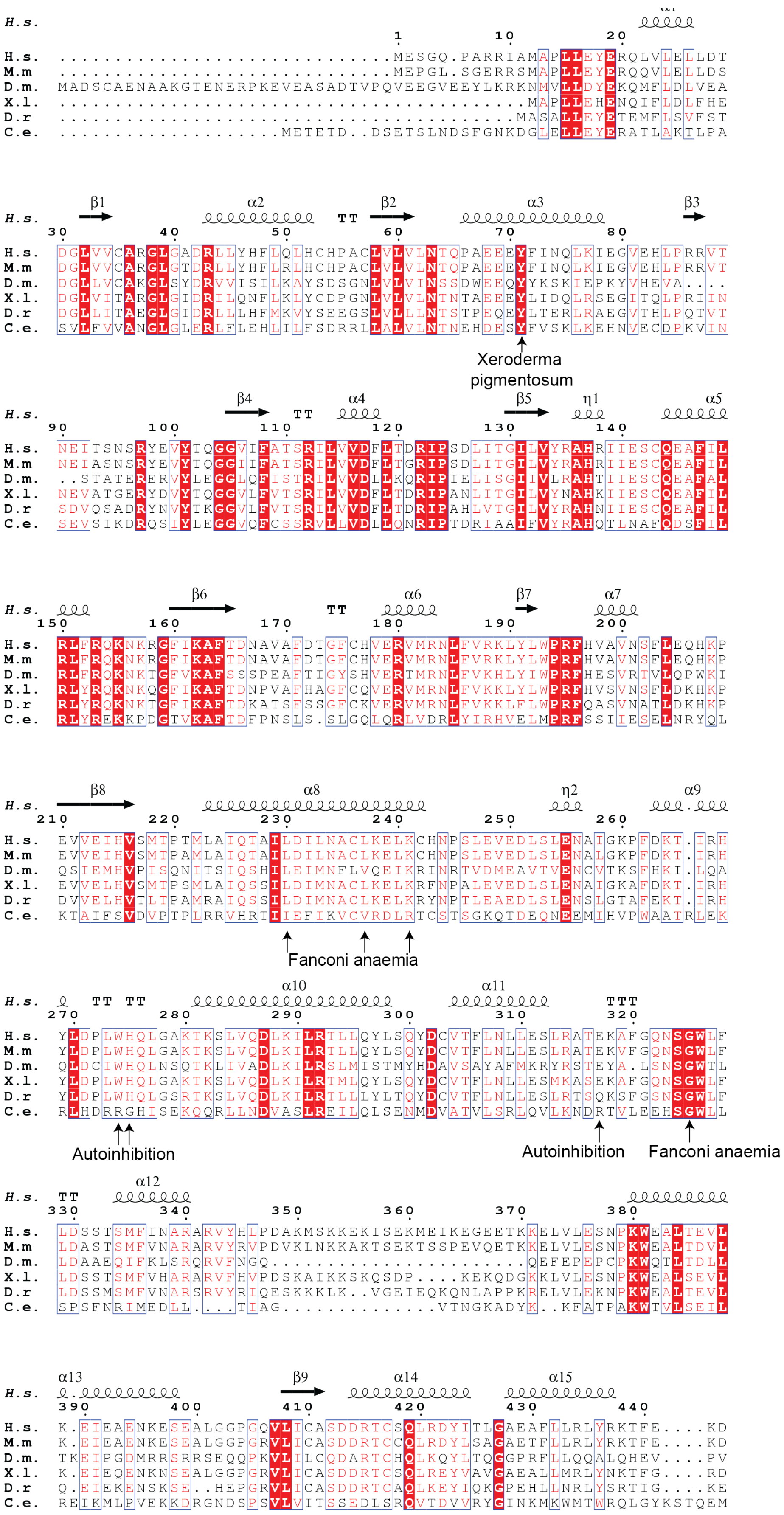
H.s.
M.m
D.m.
X.l.
D.r
C.e.
Xeroderma pigmentosum
Fanconi anaemia
Autoinhibition
Autoinhibition
Fanconi anaemia

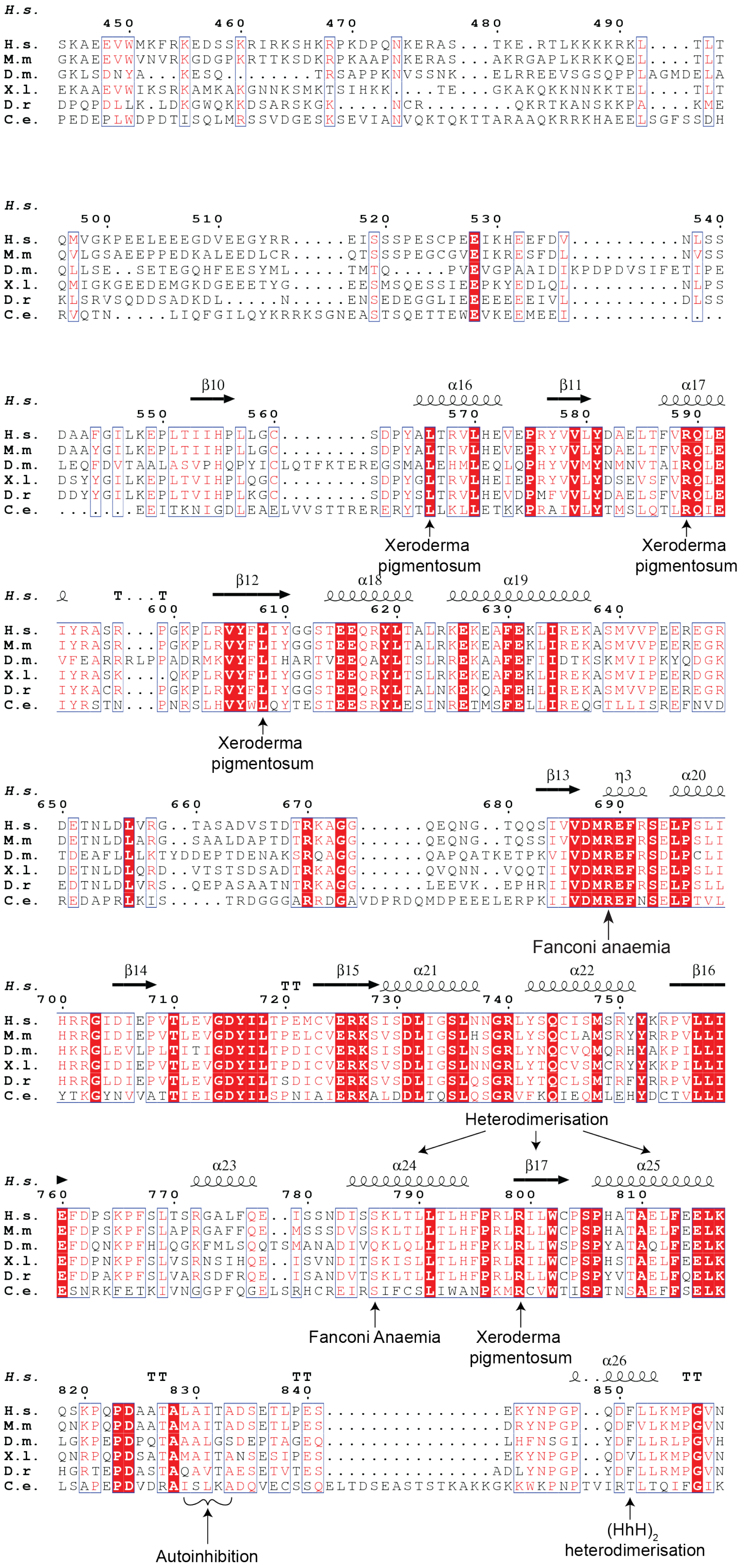
H.s. 450 460 470 480 490
H.s. SKAEEVWMKFRKEDSSKRIRKSHKRPKDPQNKERAS...TKE.RTLKKKKRKL....TLT
M.m GKAEEVWVNVRKGDGPKRTTKSDKRPKAAPNKERAS...AKRGAPLKRKKQEL....TLT
D.m. GKLSDNYA...KESQ.......TRSAPPKNVSSNK...ELRREEVSGSQPPLAGMDELA
X.l. EKAAEVWIKSRKAMKAKGNNKSMKTSIHKK....TE...GKAKQKKNNKKTEL....TLT
D.r DPQPDLLK.LDKGWQKKDSARSKGK.....NCR........QKRTKANSKKPA....KME
C.e. PEDEPLWDPDTISQLMRSSVDGESKSEVIANVQKTQKTTARAAQKRRKHAEELSGFSSDH
H.s. 500 510 520 530 540
H.s. QMVGKPEELEEEGDVEEGYRR.....EISSSPESCPEEIKHEEFDV..........NLSS
M.m QVLGSAEEPPEDKALEEDLCR.....QTSSSPEGCGVEIKREESFDL..........NVSS
D.m. QLLSE..SETEGQHFEESYML.....TMTQ.....PVEVGPAAIDIKPDPDVSIFETIPE
X.l. QMIGKGEEDEMGKDGEEETYG.....EESMSQESSIEEPKYEDLQL..........NLPS
D.r KLSRVSQDDSADKDL.....N.....ENSEDEGGLIEEEEEEEIVL..........DLSS
C.e. RVQTN.....LIQFGILQYKRRKSGNEASTSQETTEWEVKEEMEEI.............
β10 α16 β11 α17
H.s. 550 560 570 580 590
H.s. DAAFGILKEPLTIIHPLLGC........SDPYALTRVLHEVEPRYVVLYDAELTFVRQLE
M.m DAAYGILKEPLTIIHPLLGC........SDPYALTRVLHEVEPRYVVLYDAELTFVRQLE
D.m. LEQFDVTAALASVPHQPYICLQTFKTEREGSMALEHMLEQLQPHYVVMYNMNVTAIRQLE
X.l. DSYYGILKEPLTVIHPLQGC........SDPYGLTRVLHEIEPRYVVLYDSEVSFVRQLE
D.r DDYYGILKEPLTVIHPLKGC........SDPYSLTRVLHEVDPMFVVLYDAELSFVRQLE
C.e. .......EEITKNIGDLEAELVVSTTRERERYTLLKLLETKKPRAIVLYTMSLQTLRQIE
Xeroderma pigmentosum
Xeroderma pigmentosum
β12 α18 α19
H.s. T...T 600 610 620 630 640
H.s. IYRASR...PGKPLRVYFLIYGGSTEEQRYLTALRKEKEAFEKLIREKASMVVPEEREGR
M.m IYRASR...PGKPLRVYFLIYGGSTEEQRYLTALRKEKEAFEKLIREKASMVVPEEREGR
D.m. VFEARRRLPPADRMKVYFLIHARTVEEQAYLTSLRREKAAFEFIIDTKSKMVIPKYQDGK
X.l. IYRASK....QKPLRVYFLIYGGSTEEQRYLTSLRKEKEAFEKLIREKASMVIPEERDGR
D.r IYKACR...PGKPLRVYFLIYGGSTEEQRYLTALNKEKQAFEHLIREKASMVVPEEREGR
C.e. IYRSTN...PNRSLHVYWLQYTESTEESRYLESINRETMSFELLIREQGTLLISREFNVD
Xeroderma pigmentosum
β13 η3 α20
H.s. 650 660 670 680 690
H.s. DETNLDLVRG..TASADVSTDTRKAGG......QEQNG..TQQSIVVDMREFRSELPSLI
M.m DETNLDLARG..SAALDAPTDTRKAGG......QEQNG..TQSSIVVDMREFRSELPSLI
D.m. TDEAFLLKTYDDEPTDENAKSRQAGG......QAPQATKETPKVIVDMREFRSDLPCLI
X.l. DETNLDLQRD..VTSTSDSADTRKAGG......QVQNN..VQQTIIVDMREFRSELPSLI
D.r EDTNLDLVRS..QEPASAATNTRKAGG......LEEVK..EPHRIIVDMREFRSELPSLL
C.e. REDAPRLKIS.....TRDGGGARRDGAVDPRDQMDPEEELERPKIIVDMREFNSELPTVL
Fanconi anaemia
β14 TT β15 α21 α22 β16
H.s. 700 710 720 730 740 750
H.s. HRRGIDIEPVTLEVGDYILTPEMCVERKSISDLIGSLNNGRLYSQCISMSRYYKRPVLLI
M.m HRRGIDIEPVTLEVGDYILTPELCVERKSVSDLIGSLHSGRLYSQCLAMSRYYRRPVLLI
D.m. HKRGLEVLPLTITIGDYILTPDICVERKSISDLIGSLNSGRLYNQCVQMQRHYAKPILLI
X.l. HRRGIDIEPVTLEVGDYILTPDICVERKSVSDLIGSLNNGRLYTQCVSMCRYYKKPILLI
D.r HRRGLDIEPVTLEVGDYILTSDICVERKSVSDLIGSLQSGRLYTQCLSMTRFYRRPVLLI
C.e. YTKGYNVVATTIEIGDYILSPNIAIERKALDDLTQSLQSGRVFKQIEQMLEHYDCTVLLI
Heterodimerisation
α23 α24 β17 α25
H.s. 760 770 780 790 800 810
H.s. EFDPSKPFSLTSRGALFQE..ISSNDISSKLTLLTLHFPRLRILWCPSPHATAELFEELK
M.m EFDPSKPFSLAPRGAFFQE..MSSSDVSSKLTLLTLHFPRLRLLWCPSPHATAELFEELK
D.m. EFDQNKPFHLQGKFMLSQQTSMANADIVQKLQLLTLHFPKLRLIWSPSPYATAQLFEELK
X.l. EFDPNKPFSLVSRNSIHQE..ISVNDITSKISLLTLHFPRLRILWSPSPHSTAELFEELK
D.r EFDPAKPFSLVARSDFRQE..ISANDVTSKLTLLTLHFPRLRLLWCPSPYVTAELFQELK
C.e. ESNRKFETKIVNGGPFQGELSRHCREIRSIFCSLIWANPKMRCVWTISPTNSAEFFSELK
Fanconi Anaemia
Xeroderma pigmentosum
α26
H.s. TT TT TT
820 830 840 850
H.s. QSKPQPDAATALAITADSETLPES...............EKYNPGP..QDFLLKMPGVN
M.m QNKPQPDAATAMAITADSETLPES...............DRYNPGP..QDFVLKMPGVN
D.m. LGKPEPDPQTAAALGSDEPTAGEQ...............LHFNSGI..YDFLLRLPGVH
X.l. QNRPQPDSATAMAITANSESIPES...............EKYNPGP..QDVLLKMPGVN
D.r HGRTEPDASTAQAVTAESETVTES...............ADLYNPGP..YDFLLRMPGVN
C.e. LSAPEPDVDRAISLKADQVECSSQELTDSEASTSTKAKKGKKWKPNPTVIRTLTQIFGIK
Autoinhibition
(HhH)2 heterodimerisation

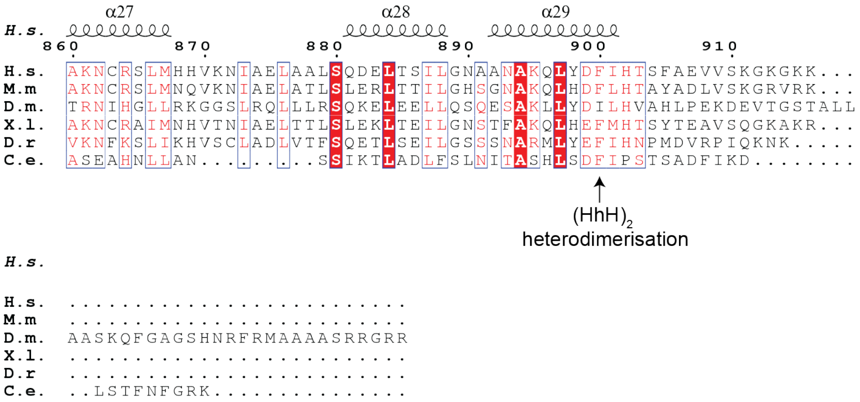

**Figure S12. Multiple sequence alignments for XPF**. *H. sapiens (H. s), Mus musculus (M. s), D. melanogaster (D. m), C. elegans (C. e), D. rerio (D. r) and X. laevis (X. l)* XPF sequences aligned with secondary structural elements above the alignment shown and annotated as used throughout the text. Residues are coloured by residue type and invariant residues indicated or highlighted by conservation.

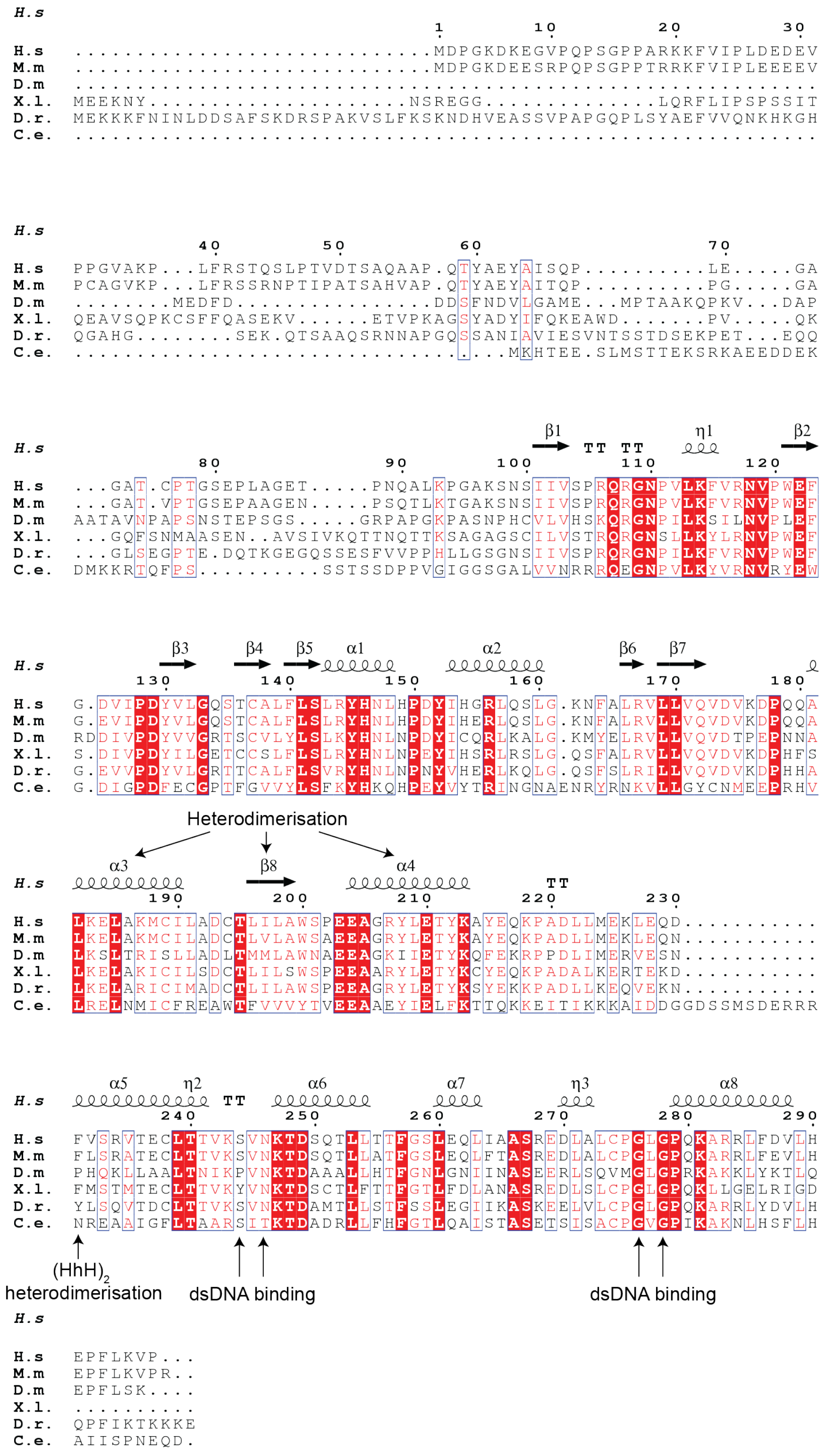


**Figure S13. Multiple sequence alignments for ERCC1.** *H. sapiens (H. s), Mus musculus (M. s), D. melanogaster (D. m), C. elegans (C. e), D. rerio (D. r) and X. laevis (X. l)* ERCC1 sequences aligned with secondary structural elements above the alignment shown and annotated as used throughout the text. Residues are coloured by residue type and invariant residues indicated or highlighted by conservation.

### SUPPLEMENTARY TABLES

| XPF residues | Wild Type Sequence | Modelled |
|---|---|---|
| 156-159 | NKRG | Poly-alanine |
| 170-172 | AFD | Poly-alanine |
| 174-177 | GFCH | Poly-alanine |
| 250-254 | EDLSL | Poly-alanine |
| 322-323 | QN | Poly-alanine |
| 345-374 | HLPDAKMSKKEKISEKMEIKEGEETKKELV | Poly-alanine |
| 396-403 | KESEALGG | Poly-alanine |
| 440-441 | FE | Poly-alanine |
| 582 | D | Poly-alanine |
| 640-641 | MV | Poly-alanine |
| 680 | T | Poly-alanine |
| 771-780 | SRGALFQEIS | Poly-alanine |
| 824-845 | DAATALAITADSETLPESEKYN | Poly-alanine |
| 441-550 | EKDSKAEEVWMKFRKEDSSKRIRKSHKRP KDPQNKERASTKERTLKKKKRKLTLTQMG KPEELEEEGDVEEGYRREISSSPESCPEEI KHEEFDVNLSSDAAFGILKEP | Not modelled |
| 642-679 | VPEEREGRDETNLDLVRGTASADVSTDTR KAGGQEQNG | Not modelled |
| 907-916 | EVVSKGKGKK | Not modelled |

| ERCC1 residues | Wild Type Sequence | Modelled |
|---|---|---|
| 223-229 | LMEKLEQ | Poly-alanine |
| 293-297 | FLKVP | Poly-alanine |
| 1-99 | MDPGKDKEGVPQPSGPPARKKFVIPLDED EVPPGVAKPLFRSTQSLPTVDTSAQAAPQT YAEYAISQPLEGAGATCPTGSEPLAGETPN QALKPGAKSN | Not modelled, evidence from SDS-PAGE that it is likely to have been proteolytically removed |

**Supplementary Table S1.** Regions in XPF and ERCC1 modelled as poly-alanine or omitted from the structure

| Id | Score | Prot1 | PP1 | PSeq1 | LP1 | Prot2 | PP2 | PSeq2 | LP2 |
|---|---|---|---|---|---|---|---|---|---|
| 3 | 50.8 | XPF | 455 | KEDSSKR | 1 | XPF | 478 | ASTKER | 4 |
| 5 | 37.34 | XPF | 438 | KTFEKDSK | 5 | XPF | 478 | ASTKER | 4 |
| 6 | 30.83 | XPF | 596 | ASRPGKPLR | 6 | XPF | 478 | ASTKER | 4 |
| 7 | 154.38 | ERCC1 | 235 | VTECLTTVKSVNK | 9 | XPF | 283 | SLVQDLKILR | 7 |
| 9 | 108.99 | XPF | 438 | KTFEKDSKAEEVWMK | 5 | XPF | 365 | EGEETKKELVLESNPK | 6 |
| 13 | 25.47 | XPF | 471 | DPQNKER | 5 | XPF | 464 | KSHKRPK | 4 |
| 15 | 36.47 | ERCC1 | 235 | VTECLTTVKSVNK | 9 | XPF | 478 | ASTKER | 4 |
| 17 | 40.04 | XPF | 625 | KEKEAFEK | 3 | XPF | 468 | RPKDPQNKER | 8 |
| 18 | 35.51 | XPF | 468 | RPKDPQNK | 3 | XPF | 626 | EKEAFEK | 2 |
| 19 | 31.84 | XPF | 596 | ASRPGKPLR | 6 | XPF | 471 | DPQNKER | 5 |
| 24 | 81.88 | XPF | 357 | ISEKMEIK | 4 | XPF | 478 | ASTKER | 4 |
| 25 | 40 | XPF | 361 | MEIKEGEETKK | 4 | XPF | 354 | KEKISEK | 3 |
| 26 | 36.47 | XPF | 371 | KELVLESNPK | 1 | XPF | 455 | KEDSSK | 1 |
| 27 | 43.77 | XPF | 357 | ISEKMEIK | 4 | XPF | 456 | EDSSKR | 5 |
| 28 | 81.46 | XPF | 355 | EKISEKMEIK | 6 | XPF | 365 | EGEETKKELVLESNPK | 6 |
| 31 | 65.29 | XPF | 343 | VYHLPDAKMSK | 8 | XPF | 455 | KEDSSKR | 1 |
| 32 | 52.59 | XPF | 855 | MPGVNAKNCR | 7 | XPF | 865 | SLMHHVK | 1 |
| 35 | 40.2 | XPF | 343 | VYHLPDAKMSKK | 8 | XPF | 455 | KEDSSKR | 6 |
| 36 | 29.69 | XPF | 343 | VYHLPDAKMSK | 8 | XPF | 455 | KEDSSKR | 5 |
| 37 | 36.47 | XPF | 365 | EGEETKKELVLESNPK | 6 | XPF | 355 | EKISEKMEIK | 2 |
| 40 | 62.69 | XPF | 343 | VYHLPDAKMSKK | 8 | XPF | 361 | MEIKEGEETKKELVLESNPK | 10 |
| 41 | 25.08 | XPF | 443 | DSKAEEVWMK | 3 | XPF | 455 | KEDSSKR | 1 |
| 42 | 49.03 | XPF | 357 | ISEKMEIK | 4 | XPF | 438 | KTFEKDSKAEEVWMK | 5 |
| 43 | 17.1 | XPF | 365 | EGEETKKELVLESNPK | 6 | XPF | 456 | EDSSKR | 5 |
| 44 | 39.97 | XPF | 443 | DSKAEEVWMK | 3 | XPF | 455 | KEDSSKR | 5 |
| 45 | 81.52 | XPF | 365 | EGEETKKELVLESNPK | 7 | XPF | 438 | KTFEKDSK | 5 |
| 46 | 33.32 | ERCC1 | 235 | VTECLTTVKSVNK | 9 | XPF | 471 | DPQNKER | 5 |
| 47 | 51.84 | XPF | 343 | VYHLPDAKMSK | 8 | XPF | 471 | DPQNKER | 5 |
| 48 | 23.75 | XPF | 343 | VYHLPDAKMSK | 10 | XPF | 471 | DPQNKER | 5 |
| 50 | 102.4 | XPF | 343 | VYHLPDAKMSKK | 8 | XPF | 354 | KEKISEK | 3 |
| 51 | 42.47 | XPF | 371 | KELVLESNPK | 1 | XPF | 354 | KEKISEKMEIK | 3 |
| 53 | 46.46 | XPF | 626 | EKEAFEK | 2 | ERCC1 | 1 | MDPGKDKEGVPQPSGPPA R | 5 |
| 54 | 32.66 | XPF | 439 | TFEKDSKAEEVWMK | 7 | XPF | 478 | ASTKER | 4 |
| 56 | 25.17 | XPF | 438 | KTFEKDSK | 1 | XPF | 365 | EGEETKKELVLESNPK | 6 |
| 57 | 76.72 | XPF | 443 | DSKAEEVWMK | 3 | XPF | 365 | EGEETKKELVLESNPK | 6 |
| 59 | 73.97 | XPF | 446 | AEEVWMKFR | 7 | XPF | 455 | KEDSSKR | 1 |
| 62 | 113.08 | XPF | 343 | VYHLPDAKMSK | 8 | XPF | 361 | MEIKEGEETKK | 4 |
| 63 | 39.97 | XPF | 636 | EKASMVVPEER | 2 | XPF | 626 | EKEAFEK | 2 |
| 65 | 131.65 | XPF | 361 | MEIKEGEETK | 4 | XPF | 371 | KELVLESNPK | 1 |
| 66 | 36.56 | XPF | 443 | DSKAEEVWMK | 3 | XPF | 355 | EKISEK | 2 |
| 68 | 60.96 | XPF | 855 | MPGVNAKNCR | 7 | ERCC1 | 1 | MDPGKDKEGVPQPSGPPA R | 5 |
| 69 | 34.81 | XPF | 361 | MEIKEGEETKK | 4 | XPF | 443 | DSKAEEVWMK | 3 |
| 71 | 78.53 | XPF | 443 | DSKAEEVWMK | 3 | XPF | 438 | KTFEK | 1 |
| 74 | 92.19 | XPF | 343 | VYHLPDAKMSKK | 8 | XPF | 365 | EGEETKKELVLESNPK | 7 |
| 75 | 33.41 | XPF | 446 | AEEVWMKFR | 7 | XPF | 455 | KEDSSKR | 5 |
| 76 | 74.85 | XPF | 365 | EGEETKKELVLESNPK | 7 | XPF | 357 | ISEKMEIKEGEETK | 4 |
| 77 | 103.29 | XPF | 343 | VYHLPDAKMSKK | 8 | XPF | 355 | EKISEKMEIK | 6 |
| 78 | 58.98 | XPF | 343 | VYHLPDAKMSKK | 10 | XPF | 355 | EKISEKMEIK | 6 |
| 79 | 57.99 | XPF | 446 | AEEVWMKFR | 7 | XPF | 478 | ASTKER | 4 |
| 83 | 114.28 | XPF | 446 | AEEVWMKFR | 7 | XPF | 365 | EGEETKKELVLESNPK | 6 |
| 84 | 77.06 | ERCC1 | 1 | MDPGKDKEGVPQPSGPPAR | 5 | ERCC1 | 235 | VTECLTTVKSVNK | 9 |
| 85 | 11.44 | XPF | 365 | EGEETKKELVLESNPK | 6 | XPF | 343 | VYHLPDAKMSK | 10 |
| 86 | 27.1 | XPF | 446 | AEEVWMKFR | 7 | XPF | 455 | KEDSSKR | 6 |
| 88 | 88.63 | ERCC1 | 235 | VTECLTTVKSVNK | 9 | XPF | 281 | TKSLVQDLK | 2 |
| 89 | 34.1 | ERCC1 | 1 | MDPGKDKEGVPQPSGPPAR | 5 | XPF | 636 | EKASMVVPEER | 2 |
| 90 | 54.44 | XPF | 446 | AEEVWMKFR | 7 | XPF | 438 | KTFEKDSK | 5 |
| 91 | 41.28 | XPF | 446 | AEEVWMKFR | 7 | XPF | 471 | DPQNKER | 5 |
| 93 | 41.5 | ERCC1 | 157 | LQSLGKNFALR | 6 | XPF | 855 | MPGVNAKNCR | 7 |
| 95 | 77.75 | ERCC1 | 214 | AYEQKPADLLMEKLEQDFVS R | 5 | XPF | 855 | MPGVNAKNCR | 7 |
| 96 | 61.32 | XPF | 443 | DSKAEEVWMK | 3 | XPF | 357 | ISEKMEIK | 4 |
| 97 | 65.83 | XPF | 443 | DSKAEEVWMKFR | 10 | XPF | 438 | KTFEK | 1 |
| 98 | 138.75 | XPF | 439 | TFEKDSKAEEVWMK | 7 | XPF | 371 | KELVLESNPK | 1 |
| 99 | 62.26 | ERCC1 | 109 | GNPVLKFVR | 6 | XPF | 626 | EKEAFEK | 2 |
| 100 | 43.81 | XPF | 343 | VYHLPDAKMSK | 8 | XPF | 443 | DSKAEEVWMK | 3 |
| 102 | 28.63 | XPF | 283 | SLVQDLKILR | 7 | XPF | 455 | KEDSSKR | 1 |
| 104 | 51.85 | ERCC1 | 269 | EDLALCPGLGPQKAR | 13 | ERCC1 | 235 | VTECLTTVKSVNK | 9 |
| 105 | 55.06 | XPF | 491 | KLTLTQMVGKPEELEEEGDVE EGYRR | 1 | XPF | 478 | ASTKER | 4 |

| | | | | | | | | | |
|---|---|---|---|---|---|---|---|---|---|
| **106** | 83.23 | ERCC1 | 109 | GNPVLKFVR | 6 | ERCC1 | 1 | MDPGKDKEGVPQPSGPPA R | 5 |
| **109** | 115.71 | XPF | 636 | EKASMVVPEER | 2 | XPF | 189 | KLYLWPR | 1 |
| **110** | 35.1 | XPF | 390 | EIEAENKESEALGGPGQVLIC ASDDR | 7 | XPF | 596 | ASRPGKPLR | 6 |
| **111** | 74.16 | ERCC1 | 1 | MDPGKDKEGVPQPSGPPAR | 5 | ERCC1 | 157 | LQSLGKNFALR | 6 |
| **112** | 35.1 | XPF | 491 | KLTLTQMVGKPEELEEEGDVE EGYRR | 1 | XPF | 596 | ASRPGKPLR | 6 |
| **113** | 69.09 | XPF | 636 | EKASMVVPEER | 2 | ERCC1 | 109 | GNPVLKFVR | 6 |
| **114** | 81.34 | ERCC1 | 214 | AYEQKPADLLMEK | 5 | ERCC1 | 1 | MDPGKDKEGVPQPSGPPA R | 5 |
| **115** | 56.56 | ERCC1 | 168 | VLLVQVDVKDPQQALK | 9 | XPF | 626 | EKEAFEK | 2 |
| **118** | 70.55 | ERCC1 | 1 | MDPGKDKEGVPQPSGPPAR | 5 | ERCC1 | 168 | VLLVQVDVKDPQQALK | 9 |
| **119** | 55.39 | XPF | 491 | KLTLTQMVGKPEELEEEGDVE EGYRR | 10 | XPF | 625 | KEKEAFEK | 3 |
| **120** | 218.79 | XPF | 671 | KAGGQEQNGTQQSIVVDMR | 1 | ERCC1 | 168 | VLLVQVDVKDPQQALK | 9 |
| **122** | 89.46 | ERCC1 | 214 | AYEQKPADLLMEK | 5 | ERCC1 | 109 | GNPVLKFVR | 6 |
| **125** | 67.98 | XPF | 242 | CHNPSLEVEDLSLENAIGKPF DKTIR | 19 | ERCC1 | 235 | VTECLTTVKSVNK | 9 |
| **128** | 76.8 | ERCC1 | 168 | VLLVQVDVKDPQQALK | 9 | ERCC1 | 109 | GNPVLKFVR | 6 |
| **129** | 106.29 | ERCC1 | 269 | EDLALCPGLGPQKAR | 13 | XPF | 283 | SLVQDLKILR | 7 |
| **131** | 41.5 | ERCC1 | 21 | KFVIPLDEDEVPPGVAKPLFR | 17 | XPF | 855 | MPGVNAKNCR | 7 |
| **133** | 25.47 | XPF | 381 | WEALTEVLKEIEAENK | 9 | XPF | 455 | KEDSSKR | 5 |
| **137** | 20.9 | XPF | 315 | ATEKAFGQNSGWLFLDSSTS MFINAR | 4 | XPF | 361 | MEIKEGEETKK | 4 |
| **138** | 20.81 | XPF | 381 | WEALTEVLKEIEAENKESEAL GGPGQVLICASDDR | 9 | XPF | 478 | ASTKER | 4 |
| **139** | 66.66 | XPF | 381 | WEALTEVLKEIEAENK | 9 | XPF | 455 | KEDSSKR | 1 |
| **141** | 52.45 | XPF | 896 | QLYDFIHTSFAEVVSKGK | 15 | ERCC1 | 214 | AYEQKPADLLMEK | 5 |
| **142** | 29.89 | XPF | 381 | WEALTEVLKEIEAENK | 9 | XPF | 455 | KEDSSKR | 6 |
| **144** | 88.94 | ERCC1 | 22 | FVIPLDEDEVPPGVAKPLFR | 16 | ERCC1 | 214 | AYEQKPADLLMEK | 5 |
| **146** | 76.85 | XPF | 751 | YYKRPVLLIEFDPSKPFSLTSR | 3 | ERCC1 | 157 | LQSLGKNFALR | 6 |
| **149** | 138.38 | ERCC1 | 214 | AYEQKPADLLMEKLEQDFVS R | 13 | XPF | 855 | MPGVNAKNCR | 7 |
| **150** | 114.5 | ERCC1 | 22 | FVIPLDEDEVPPGVAKPLFR | 16 | ERCC1 | 168 | VLLVQVDVKDPQQALK | 9 |
| **157** | 91.63 | XPF | 773 | GALFQEISSNDISSKLTLLTLH FPR | 15 | ERCC1 | 214 | AYEQKPADLLMEK | 5 |
| **161** | 45.16 | XPF | 626 | EKEAFEK | 2 | XPF | 465 | SHKRPK | 3 |
| **162** | 24.42 | XPF | 361 | MEIKEGEETKK | 4 | XPF | 439 | TFEKDSK | 4 |
| **164** | 7.98 | XPF | 371 | KELVLESNPK | 1 | XPF | 456 | EDSSKR | 5 |
| **165** | 27.95 | XPF | 365 | EGEETKKELVLESNPK | 7 | XPF | 455 | KEDSSKR | 4 |
| **167** | 20.08 | XPF | 365 | EGEETKKELVLESNPK | 6 | XPF | 471 | DPQNKER | 5 |
| **168** | 13.96 | XPF | 343 | VYHLPDAKMSK | 10 | XPF | 354 | KEKISEK | 3 |
| **170** | 31 | XPF | 443 | DSKAEEVWMK | 2 | XPF | 456 | EDSSKR | 5 |
| **171** | 30.66 | ERCC1 | 214 | AYEQKPADLLMEK | 5 | XPF | 912 | GKGKK | 2 |
| **173** | 30.44 | XPF | 443 | DSKAEEVWMK | 3 | XPF | 471 | DPQNKER | 5 |
| **175** | 45.16 | ERCC1 | 1 | MDPGKDKEGVPQPSGPPAR | 7 | XPF | 626 | EKEAFEK | 2 |
| **177** | 42.1 | XPF | 343 | VYHLPDAKMSKK | 10 | XPF | 371 | KELVLESNPK | 1 |
| **178** | 29.35 | ERCC1 | 109 | GNPVLKFVR | 6 | XPF | 438 | KTFEK | 1 |
| **183** | 31.77 | XPF | 268 | HYLDPLWHQLGAKTK | 13 | XPF | 283 | SLVQDLK | 1 |
| **186** | 12.87 | XPF | 315 | ATEKAFGQNSGWLFLDSSTS MFINAR | 4 | XPF | 355 | EKISEK | 2 |
| **187** | 32.95 | ERCC1 | 21 | KFVIPLDEDEVPPGVAKPLFR | 17 | ERCC1 | 1 | MDPGKDKEGVPQPSGPPA R | 5 |
| **192** | 17.33 | XPF | 478 | ASTKER | 4 | XPF | 438 | KTFEKDSK | 1 |
| **194** | 17.1 | ERCC1 | 1 | MDPGKDKEGVPQPSGPPAR | 5 | XPF | 478 | ASTKER | 4 |
| **195** | 23.56 | ERCC1 | 1 | MDPGKDKEGVPQPSGPPAR | 5 | XPF | 596 | ASRPGKPLR | 6 |
| **199** | 30.15 | ERCC1 | 168 | VLLVQVDVKDPQQALK | 9 | XPF | 855 | MPGVNAKNCR | 7 |
| **200** | 38.37 | XPF | 446 | AEEVWMKFR | 7 | XPF | 357 | ISEKMEIK | 4 |
| **212** | 31.84 | XPF | 381 | WEALTEVLKEIEAENK | 9 | XPF | 471 | DPQNKER | 5 |
| **215** | 40.07 | XPF | 355 | EKISEK | 2 | XPF | 478 | ASTKER | 4 |
| **226** | 18.68 | XPF | 468 | RPKDPQNK | 3 | XPF | 455 | KEDSSKR | 6 |
| **233** | 22.47 | XPF | 343 | VYHLPDAKMSK | 10 | XPF | 354 | KEKISEK | 1 |
| **238** | 18.44 | XPF | 446 | AEEVWMKFR | 7 | XPF | 465 | SHKRPKDPQNK | 3 |
| **239** | 51.69 | XPF | 446 | AEEVWMKFR | 7 | XPF | 468 | RPKDPQNKER | 3 |
| **241** | 27.65 | XPF | 491 | KLTLTQMVGKPEELEEEGDVE EGYRR | 1 | XPF | 626 | EKEAFEK | 2 |
| **245** | 27.03 | XPF | 491 | KLTLTQMVGKPEELEEEGDVE EGYRR | 1 | XPF | 438 | KTFEK | 1 |
| **248** | 71.02 | ERCC1 | 1 | MDPGKDKEGVPQPSGPPAR | 5 | XPF | 268 | HYLDPLWHQLGAKTK | 13 |
| **258** | 24.3 | XPF | 381 | WEALTEVLKEIEAENK | 9 | XPF | 465 | SHKRPK | 3 |
| **262** | 154.38 | XPF | 242 | CHNPSLEVEDLSLENAIGKPF | 19 | XPF | 283 | SLVQDLKILR | 7 |

| | | | | | | | | | |
|---|---|---|---|---|---|---|---|---|---|
| | | | | DKTIR | | | | | |
| **265** | 18.31 | XPF | 381 | WEALTEVLKEIEAENK | 5 | XPF | 455 | KEDSSK | 1 |
| **273** | 80.24 | XPF | 773 | GALFQEISSNDISSKLTLLTLH FPR | 15 | XPF | 855 | MPGVNAKNCR | 7 |
| **277** | 40.07 | XPF | 365 | EGEETKKELVLESNPK | 6 | XPF | 455 | KEDSSK | 1 |
| **280** | 12.63 | XPF | 343 | VYHLPDAKMSK | 8 | XPF | 354 | KEKISEK | 1 |
| **286** | 22.2 | XPF | 390 | EIEAENKESEALGGPGQVLIC ASDDR | 7 | XPF | 468 | RPKDPQNK | 3 |
| **293** | 12.63 | XPF | 751 | YYKRPVLLIEFDPSKPFSLTSR | 3 | XPF | 855 | MPGVNAKNCR | 7 |

**Supplementary Table S2. Mass spectrometry crosslinking data.** All verified XL-MS crosslinks displayed. Id – unique identifier, score – normalised prevalence value representing the frequency of crosslink appearance in MS2, Prot1/2 – protein from which the respective crosslink originates, PP1/2 – position in sequence for respective crosslink, Pseq1/2 – protein sequence identified in MS1, LP1/2 – link position identifying which residue in Pseq the crosslink maps to.
